# Stromal Amyloid β drives Neutrophil extracellular trap formation to augment tumour growth

**DOI:** 10.1101/2020.01.10.901686

**Authors:** Hafsa Munir, James O. Jones, Tobias Janowitz, Carla P. Martins, Sarah J. Welsh, Jacqueline D. Shields

## Abstract

Tumors are comprised of cancer cells and a network of non-cancerous stromal cells. Cancer-associated fibroblasts (CAFs) are well known to support tumorigenesis and are emerging as immune modulators. While many leukocyte populations are well studied in cancer, neutrophils have received less attention. Neutrophils can release histone-bound nuclear DNA and cytotoxic granules as extracellular traps (NETs) in a process termed NETosis. Here, we show that CAFs induce formation of NETs both within the tumor microenvironment and at systemic levels in the blood and bone marrow. These tumor-induced NETs (t-NETs) are driven by a ROS-mediated pathway dependent on PAD4 and CD11b. Remarkably, CAF-derived Amyloid β was identified as the key factor driving t-NETosis, a protein with significance in both neurodegenerative and inflammatory disorders. Therapeutic inhibition of NETs in established tumors prevented growth, skewing neutrophils to a pro-inflammatory phenotype. Reciprocally, t-NETs enhanced CAF activation phenotypes. Mirroring murine observations, NETs were detected juxtaposed to CAFs in human melanoma and pancreatic adenocarcinoma, and elevated expression of amyloid and β-Secretase correlated with poor prognosis. In summary, we report the existence of cross-talk between CAFs and neutrophils within the tumour microenvironment whereby CAF-induced t-NETosis supports cancer progression, identifying Amyloid β as the protagonist and potential therapeutic target.

**Significance:** This study defines the existence of a pro-tumor immunomodulatory function of the stroma showing the induction of Neutrophil Extracellular Traps through CAF-derived Amyloid β. We term this novel process “Tumor-induced NETosis” (t-NETosis) and propose that therapeutic inhibition of this mechanism, which we observe in human melanoma and pancreatic cancer, has the potential to improve patient outcome.

## Introduction

The tumor microenvironment comprises a complex niche of cancer cells and “normal” cell populations collectively referred to as the stroma. The stroma includes leukocytes, cancer-associated fibroblasts (CAFs), pericytes, blood vessels and lymphatic vessels ^1^. Tumor development is accompanied by changes in the phenotype, function and interactions between these stromal constituents ^2–5^, a process which is central to carcinogenesis.

CAFs are one of the most abundant stromal populations in the tumor, and display considerable heterogeneity and plasticity ^6^. Multiple tumour promoting functions have been attributed to CAFs, including promoting angiogenesis, remodelling extracellular matrix ^7,8^, modifying tumor stiffness ^9,10^, nutrient processing ^11^ and facilitating the invasion of tumor cells ^1,4^. More recently, CAFs have emerged as modulators of the innate and adaptive immune responses as they are recruited to the tumor. CAF-derived factors have been shown to drive the recruitment and M2 polarization of macrophages ^12^. CAFs drive T-cell deletion and exhaustion in an antigen-dependent manner by FASL and PD-L2 mediated interactions ^5^. CAFs also mediate exclusion of T-cells from tumors via production of CXCL12 ^2^. Furthermore, CAF-derived IL-6 can induce systemic immunosuppressive effects ^13^. Collectively, these data demonstrate that anti-cancer immune responses can be modulated by CAFs, and failure of the immune system to control cancer is not solely mediated by cancer cells but also the surrounding stroma.

Neutrophils are the most abundant circulating leukocyte population, functioning as early responders to inflammatory insult ^14^. Following activation, neutrophils utilize several mechanisms to exert their effects, such as secreting inflammatory factors that influence other immune populations, production of reactive oxygen species (ROS) and cytotoxic granular proteins to eliminate pathogens, as well as releasing extracellular traps (NETs). NETs are composed of chromatin bound DNA decorated in cytosolic and granular proteins such as myeloperoxidase (MPO) and neutrophil elastase (NE) ^14^. Though the molecular mechanisms governing NET release are still not completely understood, in certain contexts it requires ROS-mediated, calcium driven citrullination of histones by Protein Arginine Deiminase 4 (PAD4) ^14^.

NETs detected in chronic inflammatory disorders exert their pro-inflammatory effects by modulating the activity of other stromal populations at the site of tissue damage. In murine models of inflammation, such as systemic lupus erythematosus, NETs activate plasmacytoid dendritic cells through engagement of TLR9, thus exacerbating the condition ^15,16^. In addition, NETs have been shown to reduce the threshold for the activation of CD4^+^ T-cells in response to antigen ^17^. As well as regulating immune cells, NET-derived components were found to induce activation of lung fibroblasts in a lung fibrosis model, promoting their differentiation into myofibroblasts ^18^. Subsequent collagen deposition, proliferation and migration of the differentiated myofibroblasts was also enhanced by treatment with NETs ^18^.

While the roles of neutrophils in infection have been well established, their contribution to tumor progression, immune evasion and metastasis remains controversial. Indeed, neutrophils have been reported to exert both anti- and pro-tumorigenic effects depending on the environmental cues to which they are exposed ^19,20^. This plasticity has made it difficult to ascertain the spatial and temporal nature of neutrophil functions within tumors. Neutrophil-derived NETs have been identified as modulators of cancer-induced thrombosis through a granulocyte-colony stimulating factor (G-CSF) dependent mechanism ^21^ and facilitate metastasis by capturing circulating tumor cells to promote colonization of distal sites ^22–25^. They have also been detected in several blood cancers ^26–28^ and recently, NET-mediated remodelling of the extracellular matrix has been reported to awaken dormant cancer cells and promote aggressive lung metastasis ^29^. While observed in primary tumors, the contribution of NETs to tumorigenesis and underlying mechanisms are lacking ^22,30–34^.

Here, we report the existence of a previously undescribed crosstalk between CAFs and neutrophils driving a phenomenon termed Tumor-induced NETosis (t-NETosis). In three different models of cancer we show that CAF-secreted Amyloid β drives formation of t-NETs through CD11b in a ROS-dependent, PAD4-driven mechanism both within the microenvironment and at systemic levels in the blood and bone marrow. Therapeutic inhibition of PAD4 stopping t-NETosis, or prevention of Amyloid β release abolished growth of established tumors and restored a pro-inflammatory-like status indicating a potential axis to be exploited therapeutically.

## Results

While neutrophils are observed in tumors, little is known about how micro-niches created by stromal compartments perturb the activity of these immune cells. In primary murine pancreatic and skin (melanoma) tumors we observed that neutrophils were frequently confined to CAF-rich regions (Figure 1A) implying a potential for cross-talk between the two populations.

**Fig. 1.**
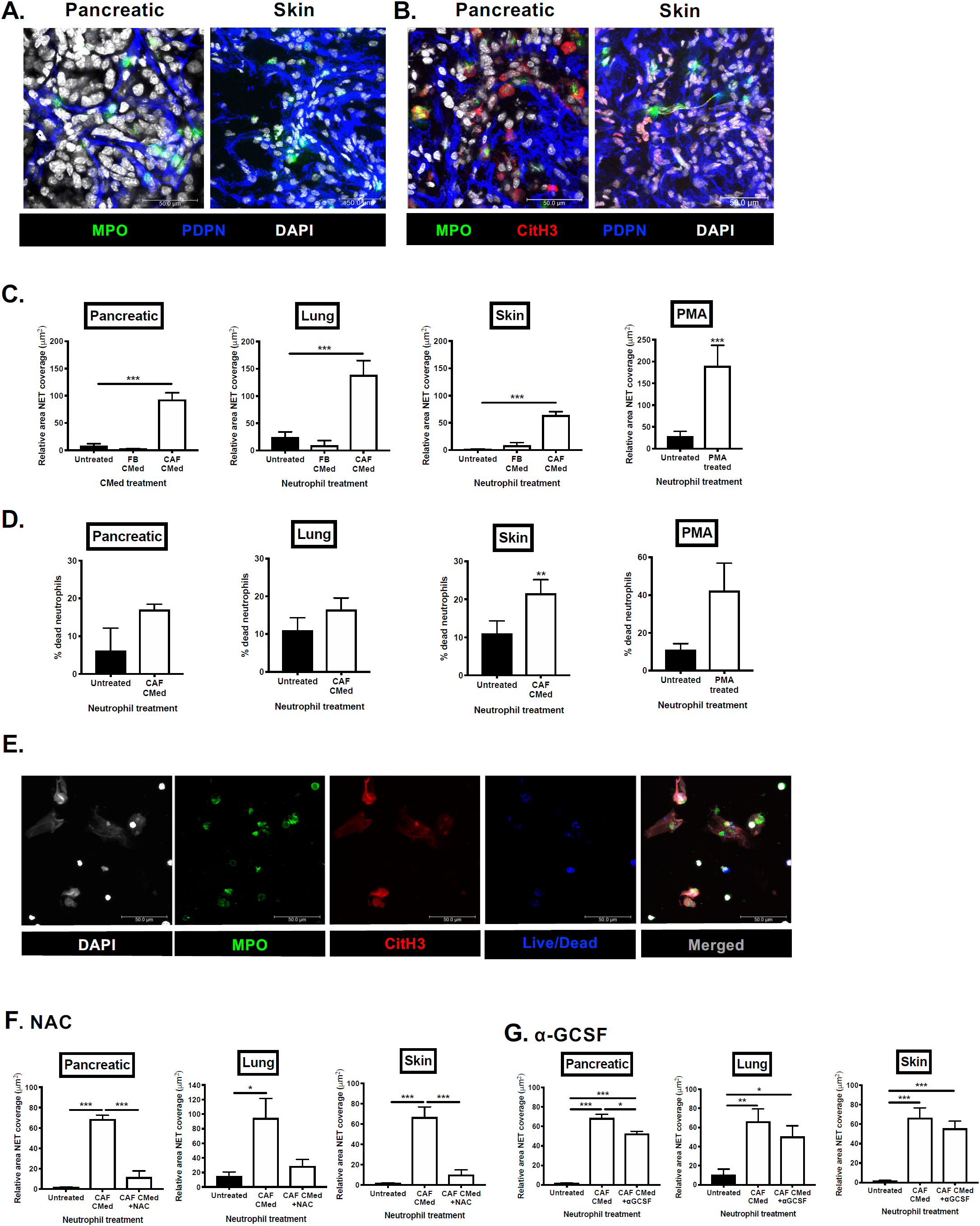
CAF-derived factors induce NETosis *in vitro* and *in vivo*. (**A**) Confocal microscopy of myeloperoxidase (MPO) and podoplanin (PDPN) expressed on neutrophils and CAFs respectively in pancreatic and skin tumors. (**B**) Confocal microscopy of NETs in murine pancreatic and skin tumors showing expression of myeloperoxidase (MPO) and Citrullinated histone H3 (CitH3) by NETting neutrophils and podoplanin (PDPN) by CAFs. (**C**) Quantification of the area of the field covered by SYTOX green positive neutrophil-derived extracellular DNA relative to the number of neutrophils in each field after treatment with pancreatic, lung or skin FBs CMed, CAF CMed or PMA for 3h. (**D**) The percentage of dead neutrophils after treatment with pancreatic, lung or skin CAF CMed or PMA based on the number of SYTOX green positive neutrophils. (**E**) Confocal microscopy of bone marrow neutrophils stained with MPO, CitH3 and live/dead cell viability dye after induction of NETosis by treatment with CAF CMed for 3h. Quantification of the relative NET coverage of neutrophils treated with CAF CMed with or without pre-treatment with (**F**) N-acetyl-cysteine (NAC) or (**G**) anti-granulocyte colony stimulating factor (α-GCSF). Data are mean ± SEM; * = p<0.05, ** = p<0.01 and *** = p<0.001 using t-test. Assays were performed on (C) n=3-10, (D) n=4, (F) n=4-5 and (G) n=4 independent experiments. Scale bars are 50µm.

### Influence of CAF-derived factors on neutrophil function

To determine if CAF-derived factors have the capacity to impact aspects of neutrophil behavior, we first treated bone marrow-derived neutrophils with conditioned media (CMed) from CAFs, tissue-matched normal FBs, or Phorbol 12-myristate 13-acetate (PMA; a well-known activator of neutrophils) and assessed viability, surface activation markers, ROS production and phagocytic capability of the neutrophils. Isolated CAFs were characterised based on surface expression of classic markers; Thy1, Podoplanin and PDGFRα (Supplementary figure 1A) and lack of immune (CD45) and epithelial (EpCAM) markers (Supplementary Figure 1A). Lung and pancreatic FB or CAF CMed had no impact on neutrophil viability for the duration of treatment while PMA induced significant neutrophil death (Supplementary Figure 1B). Expression of activation markers (CD11b and CD18) tended to increase in response to lung CAF CMed treatment (Supplementary Figure 1C) in the presence or absence of an additional inflammatory insult (Lipopolysaccharide; LPS). CAF CMed failed to induce ROS production in neutrophils (Supplementary Figure 1D) nor did the cells exhibit enhanced phagocytic capabilities (Supplementary Figure 1E) relative to PMA after 30min treatment. Therefore, the classical functions of neutrophils ^14^ were largely unaltered by treatment with CAF-derived factors *in vitro*.

### CAF-derived factors induce NETs in primary tumors

Having already observed neutrophils in proximity to CAF-rich regions, we also detected the presence of NETs within primary murine pancreatic and skin tumors (Figure 1B). NETs were defined as staining positive for extracellular DNA, MPO and Citrullinated histone H3 (CitH3). We termed these structures tumor-induced NETs (t-NETs), and thus sought to examine the role of t-NETs in the primary tumor, focusing on the mechanisms driving their release.

To determine if CAF-derived factors drive generation of t-NETs, we treated isolated bone marrow neutrophils with CAF or FB CMed and analyzed NETosis (Supplementary Figure 2A, B and Supplementary Movie 1). While CMed from normal FBs was unable to induce NETs, CAF CMed from pancreatic, lung and skin tumors was sufficient to induce NETosis to levels comparable with PMA (Figure 1C and Supplementary figure 2B). As NETosis is thought to be largely a “suicidal” process ^14^, we next quantified neutrophil death following longer-term exposure to CAF CMed (compared to 30min treatment in Supplementary Figure 1B). After 3h of treatment with CAF CMed, a trend of increasing cell death was observed compared to untreated neutrophils and to levels comparable with PMA (Figure 1D). This was confirmed by staining of NETting neutrophils with live/dead dye (Figure 1E) indicating that CAF-influenced NETosis is a cell-death dependent effect. As autophagy has also been implicated as a mode of death in NETosis ^34^, we then quantified the level of NETosis in the presence of chloroquine, an inhibitor of autophagy. Chloroquine did not rescue the neutrophils from undergoing NETosis (Supplementary figure 2C) ruling out autophagy as a mechanism underlying CAF-induced t-NET formation.

G-CSF and intracellular ROS have been reported to induce citrullination of Histone H3 which is required for NET formation ^35^. Thus, to determine if CAF-derived factors stimulate t-NETs in a ROS-dependent manner, we first inhibited ROS production. Pre-treatment of neutrophils with N-acetylcysteine (NAC; Figure 1F) Diphenyleneiodonium (DPI), Vitamin C or Trolox (Supplementary figure 2D-F) prior to CAF CMed treatment suppressed NET formation, indicating that ROS production by neutrophils in response to CAF-derived factors is a key mediator in the process. However, unlike previous reports which showed that tumor-derived G-CSF drives NETosis systemically, here CAF-mediated NETosis was not driven by G-CSF as NET release was not significantly reduced following neutralization of G-CSF *in vitro* (Figure 1G). Together, these data suggest that CAFs are key drivers of ROS-dependent, suicidal t-NETosis and the potential factors that induce t-NETs in this context are likely distinct from those previously reported ^21,34^.

### CAF-derived factors induce systemic effects on neutrophils

We next investigated whether CAF-derived factors could render circulating neutrophils more susceptible to NETosis before being recruited into the tumor. Indeed, we observed that bone marrow-derived neutrophils isolated from pancreatic, lung and skin tumor-bearing mice displayed a greater propensity to generate t-NETs in the absence of an additional stimulus compared to neutrophils from non-tumor bearing mice (Figure 2A). With pancreatic tumors, but not for either lung or skin, this was accompanied by an increase in neutrophil death in the bone marrow (Supplementary Figure 3A). However, there was not a significant increase in the number of neutrophils isolated from tumor bearing mice (Figure 2B).

**Fig. 2.**
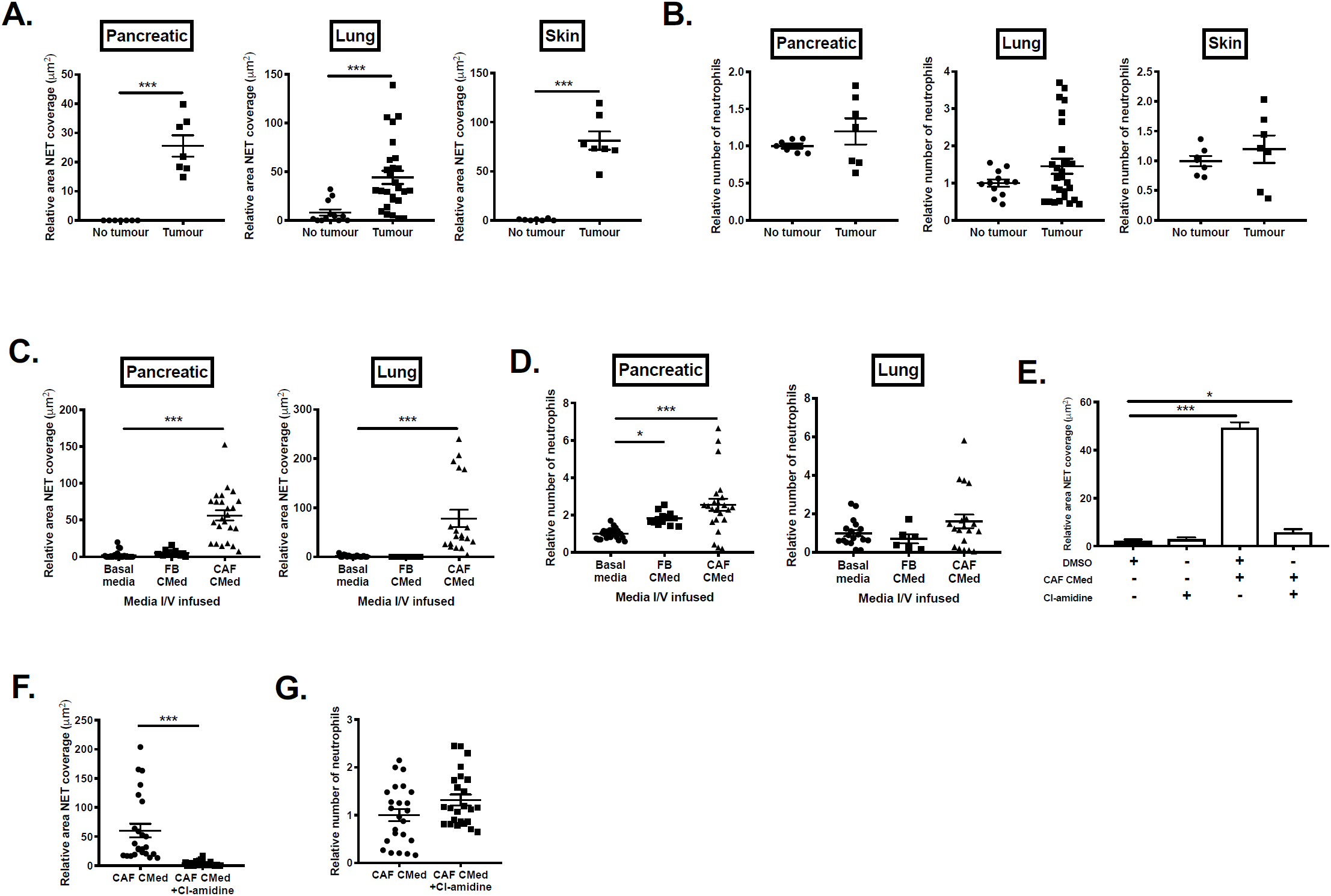
CAF-derived factors drive NETosis systemically. Quantification of (A) relative NET coverage and (B) relative number of bone marrow neutrophils taken from mice with pancreatic, lung or skin tumors. Quantification of the (C) relative NET coverage and (D) relative number of bone marrow neutrophils isolated from wild type mice intravenously infused with pancreatic or lung FB or CAF CMed 24h before analysing NETosis. (E) Quantification of relative NET coverage after stimulation with pancreatic CAF CMed with or without treatment with Cl-amidine *in vitro*. (F) Quantification of the relative NET coverage and (G) relative number of bone marrow neutrophils isolated from mice intravenously infused with lung CAF CMed with or without pre-treatment with Cl-amidine for 24h. Data are mean ± SEM; * = p<0.05, ** = p<0.01 and *** = p<0.001 using (A and F) t-test and (C-E) one-way ANOVA with a Dunnett post hoc test. Assays were performed on (A-B) n=4 (in duplicate), n=7 (in quadruplet) and n=7 for pancreatic, lung and skin tumor bearing mice respectively, (C-D) n=8 (in triplicate) and n=6-16 (in duplicate) for pancreatic and lung FB and CAF CMed infused mice respectively, (E) n=3 (in triplicate) and (F-G) n=8 (in triplicate) independent experiments.

To determine whether CAF-derived factors were sufficient to drive the observed susceptibility of bone marrow-derived neutrophils towards NETosis, we intravenously (I/V) infused CMed from pancreatic or lung-derived FBs or CAFs in the absence of tumors. Spontaneous NET production by bone marrow-derived neutrophils *ex vivo* was significantly enhanced in mice treated with CAF CMed compared to FB CMed or basal media (Figure 2C) with concurrent increases in neutrophil death (Supplementary Figure 3B) and counts (Figure 2D). Such increases in neutrophil number following CAF and FB CMed infusion indicates that FB and CAFs may be a source of other factors, such as G-CSF, in tumor bearing mice, which would be consistent with a previous report showing increases in neutrophil number in patients due to higher levels of G-CSF ^21^. The discrepancy between the neutrophil counts after I/V infusion of the CMed compared to those in the tumor bearing mice is potentially because the concentration of factors that drive neutrophil expansion may be higher in the *in vitro* CAF cultures compared to the amount secreted by the tumor stroma. Together, these data suggest that CAFs can skew neutrophils towards NET formation before they enter the tumor.

### CAF-driven t-NETs are pro-tumorigenic

Having shown that CAFs can promote t-NETosis at local and systemic levels we next sought to determine the functional impact of the t-NETs on primary tumor development. Previous studies have analyzed the effect of neutrophil depletion on tumor progression, primarily by using anti-Ly6G antibodies ^36–38^, however when studying NETosis this approach would have confounding effects as a result of their depletion. Therefore, to disrupt NETosis without impacting other neutrophil functions we inhibited PAD4, a key component of the NET pathway, driving citrullination of histones to facilitate DNA release ^14^. To first ascertain the requirement of PAD4 in CAF-driven NETosis, we tested the effects of its inhibition on neutrophils *in vitro*. A complete suppression of CAF CMed-induced NET release was observed in neutrophils treated with the PAD4 inhibitor, Cl-amidine, *in vitro* (Figure 2E). Moreover, treatment with Cl-amidine *in vivo* prior to I/V infusion of lung CAF CMed was sufficient to abolish NET release by bone marrow-derived neutrophils along with a reduction in neutrophil death *ex vivo* (Figure 2F and Supplementary Figure 3C). The number of bone marrow neutrophils was largely unaffected (Figure 2G) suggesting that NETosis was inhibited without influencing other factors driving an increase in neutrophil number.

*In vivo*, mice with established skin tumors were treated with GSK484, Cl-amidine (PAD4 inhibitors) or DMSO for 7 days (Figure 3A). Cl-amidine (Supplementary Fig 4A) treatment completely inhibited tumor growth compared with vehicle controls and this effect was replicated by treatment with the more specific PAD4 inhibitor, GSK484 (Figure 3B). While tumor volumes were drastically different following inhibition of t-NETosis, we did not observe a significant effect on tumor immune infiltrates and resident stroma by flow cytometry profiling (Figure 3C and supplementary figure 4B, C and E). Importantly, the number of neutrophils recruited to tumors was unaffected by PAD4 inhibition indicating that treatment effects on neutrophil behavior were specific to NETosis (Figure 3C, D and supplementary figure 4B). Suppression of tumor growth was not due to direct toxicity of the small molecule inhibitors on cancer cells. Indeed, treatment with GSK48, had no effect on the growth of melanoma cells *in vitro* (Supplementary figure 4F), while treatment with Cl-amidine only mildly affected cell growth (non-significant; Supplementary figure 4F) which could be a potential consequence of the broader action of Cl-amidine. Therefore, the effect of inhibiting PAD4 on the tumor growth *in vivo* was primarilty due to inhibition of NETosis.

**Fig. 3.**
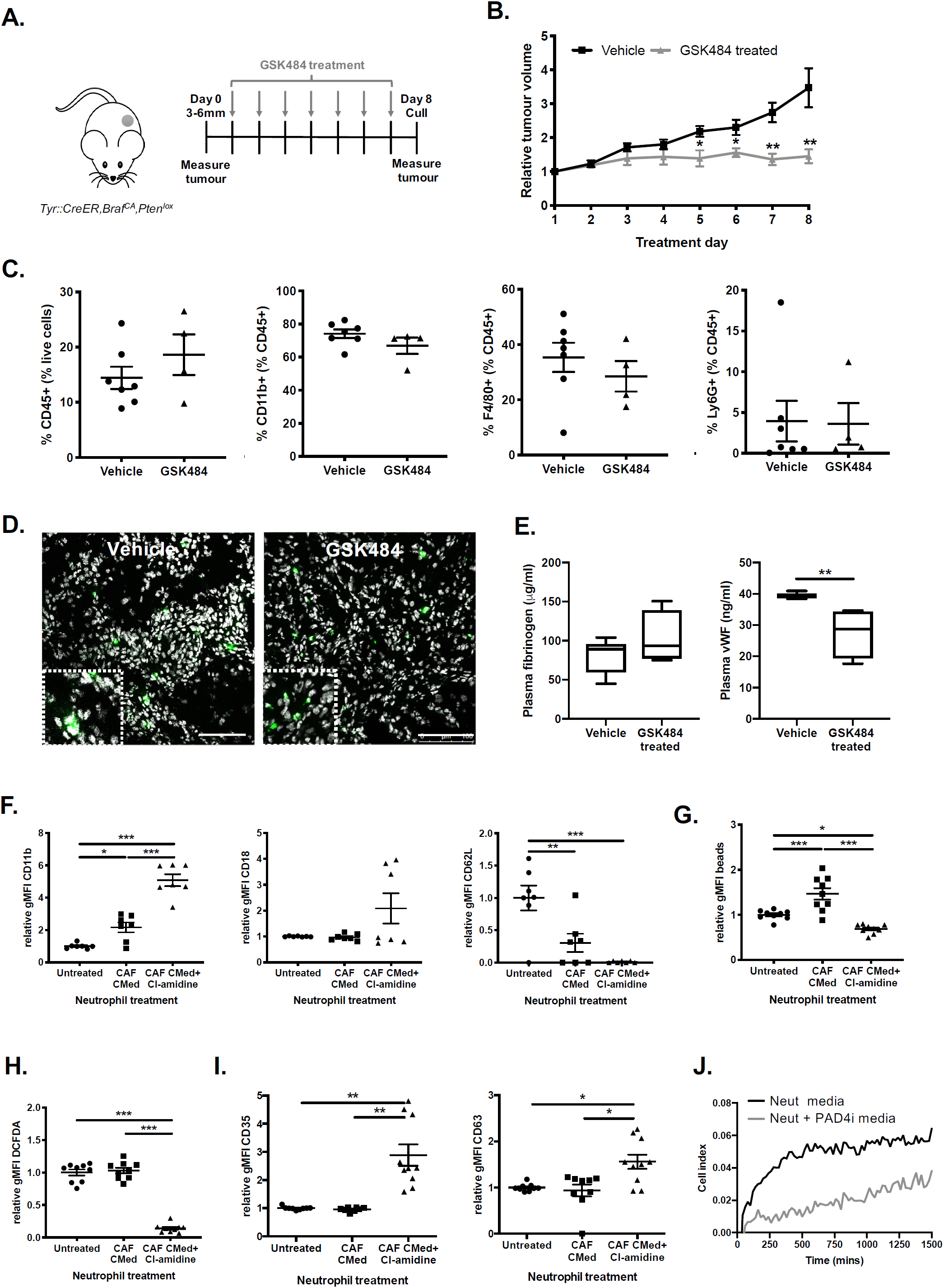
Inhibiting t-NETosis stops tumor growth in vivo. (A) Schematic of GSK484 treatment regime of skin tumor bearing mice. (B) Relative volume of skin tumors on mice treated with vehicle or GSK484 over 8d. (C) Flow cytometric analysis of the percentage immune cells (CD45^+^), myeloid cells (CD11b^+^), macrophages (F4/80^+^) and neutrophils (Ly6G^+^) in the tumor. (D) Representative confocal images showing neutrophils (myeloperoxidase (MPO) in green) within the tumor in control or treated animals (zoomed inset). Nuclei counterstained with DAPI (white). (E) The levels of clotting factors (fibrinogen and von Willebrand factor; vWF) in the plasma of skin tumor bearing mice treated with vehicle or GSK484. (F) Quantification of CD11b, CD18 and CD62L expression on wild type bone marrow-derived neutrophils after 3h treatment with pancreatic CAF CMed, with and without Cl-amidine, *in vitro* by flow cytometry. (G) The phagocytic capacity of wild type bone marrow-derived neutrophils after 3h pancreatic CAF CMed, with and without Cl-amidine, treatment based on uptake of fluorescent 1µm beads *in vitro* assessed by flow cytometry. (H) Quantification of ROS production by wild type bone marrow-derived neutrophils after 3h treatment with pancreatic CAF CMed, with and without Cl-amidine, *in vitro* based on the levels of DCFDA by flow cytometry. (I) Quantification of neutrophil degranulation based on CD35 and CD63 expression by flow cytometry after 3h pancreatic CAF CMed, with and without Cl-amidine for 3h. (J) Representative plot illustrating tumor cell growth following treatment with PAD4i-treated neutrophil derived media. Data are mean ± SEM; * = p<0.05, ** = p<0.01and *** = p<0.001 using (B and E) a Mann-Whitney test and (F-I) a one-way ANOVA with a Tukey post hoc test. Assays were performed on (B) n=7-8, (C and E) n=4-7, (F-I) n=3 (in triplicate) and (J) Representative of n=2 (in duplicate) independent experiments. Scale bars are 100µm.

**Fig. 4.**
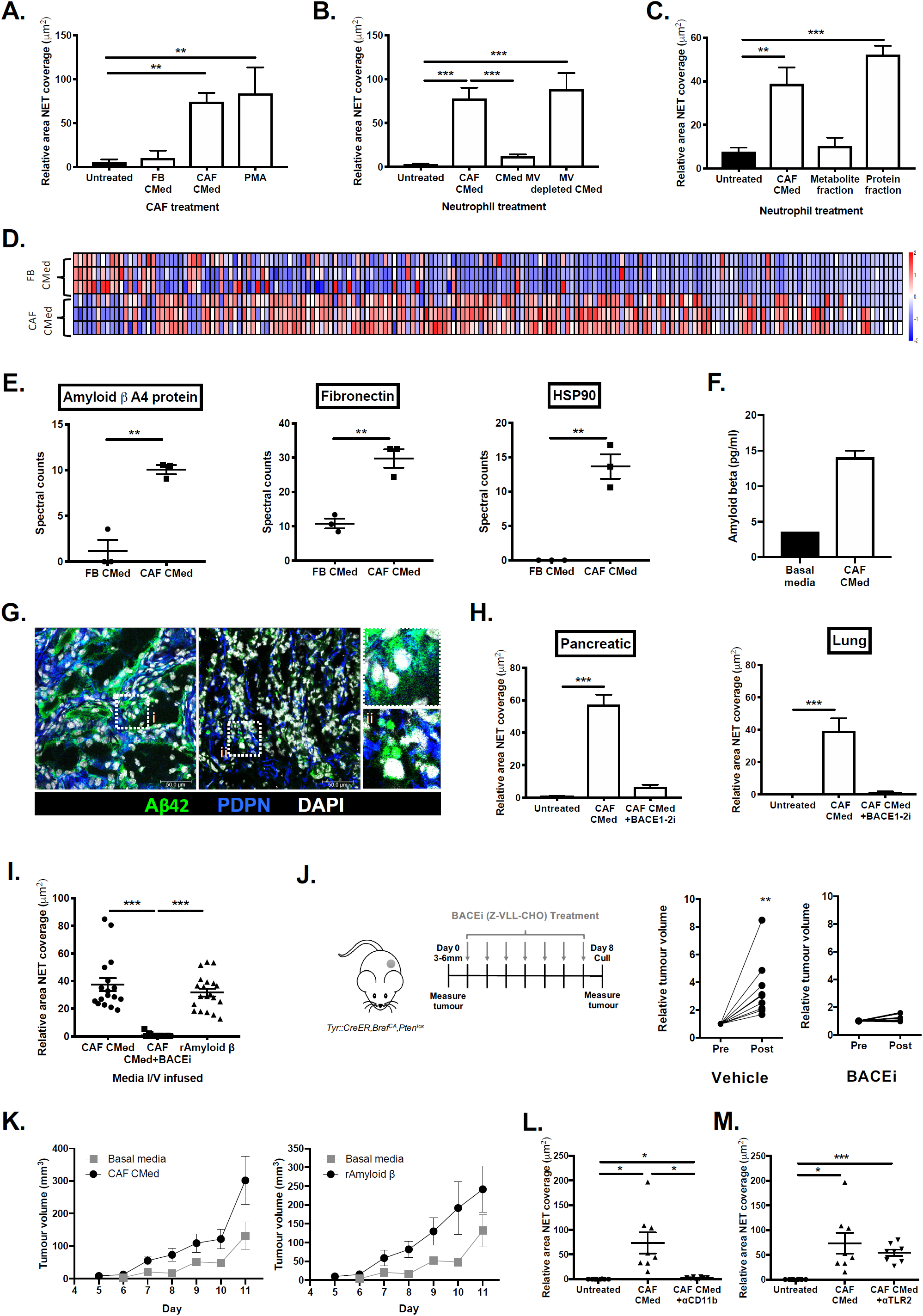
Amyloid β is the driver of CAF-induced t-NETosis. (**A**) Quantification of the relative NET coverage of neutrophils that were added to lung CAFs treated with or without FB CMed, CAF CMed or PMA for 3h. (**B**) Quantification of the relative NET coverage of neutrophils treated with lung CAF CMed or CAF CMed-derived microvesicles (MV) or CMed depleted of MV. (**C**) Quantification of the relative NET coverage of neutrophils treated with lung CAF CMed or the metabolite or protein fractions of the CAF CMed. (**D**) Differentially secreted proteins in pancreatic FB and CAF CMed analyzed by mass spectrometry. (**E**) Spectral counts for NET-related factors secreted by pancreatic FBs and CAFs. (**F**) Levels of Amyloid β in basal media and pancreatic CAF CMed. (**G**) Confocal microscopy of amyloid precursor protein (APP) and Podoplanin (PDPN) expressed by CAFs in pancreatic tumors. Insets depict diffuse vs. aggregated patterns of APP/Amyloid β distribution. (**H**) Quantification of the relative NET coverage of neutrophils treated with pancreatic and lung CAF CMed generated with or without 24h pre-treatment of the CAFs with a β-secretase 1 and 2 (BACE1-2) inhibitor. (**I**) Quantification of the relative NET coverage of bone marrow neutrophils taken from wild type mice intravenously infused with pancreatic CAF CMed taken from cells treated with or without BACE1-2 inhibitor for 24h or recombinant Amyloid β. (**J**) Schematic of BACEi treatment regime of skin tumor bearing mice and relative volume of skin tumors on mice before and after treatment with vehicle or Z-VLL-CHO. (**K**) Growth of orthotopically implanted B16.F10 tumor cells with vehicle, CAF CMed or recombinant Amyloid β treatment. Quantification of the relative NET coverage after 3h treatment with pancreatic CAF CMed with or without (**L**) CD11b or (**M**) TLR2 blocking antibodies. Data are mean ± SEM; ** = p<0.01 and *** = p<0.001 using (A and H) one-way ANOVA with a Dunnett post hoc test, (B-C, E, I and J) t-test and (L-M) one-way ANOVA with a Tukey post hoc test. Assays were performed on (A) n=3-6, (B) n=7, (C) n=5, (E) n=3, (F) n=1-3, (H) n=6 (in triplicate), (I) n=7 (in duplicate or triplicate), (J) n=7-9, (K) n=6 and (L-M) n=3 (in triplicate) independent experiments. Scale bars are 50µm.

With the knowledge that circulating neutrophil-derived NETs have been reported to contribute to thrombus formation in advanced disease we also examined the plasma of mice treated with PAD4i. GSK484, but not Cl-amidine treatment was accompanied by lower levels of the clotting factors von willebrand factor (vWF), while the levels of fibrinogen were unchanged (Figure 3E and Supplementary figure 4D) suggestive of a reduction in NET-mediated thrombosis within the circulation of treated mice.

Since the anti-tumor effects of t-NET inhibition were not directed by secondary effects on other infiltrating immune populations, we then examined the neutrophils themselves in more detail. Following treatment with CAF CMed and PAD4 inhibitor for 3h *in vitro* to prevent NETs, neutrophils increased expression of activation markers CD11b and CD18, and completely shed surface CD62L (Figure 3F), typical of activated neutrophils. Phagocytosis was reduced and intracellular ROS levels were exhausted, consistent with depletion following an oxidative burst (Figure 3G and H). Furthermore, neutrophils exhibited enhanced degranulation with increasing expression of surface CD35 and CD63 after PAD4 inhibition associated with a pro-inflammatory phenotype (Figure 3I). To test the pro-inflammatory, anti-tumour potential of degranulation following PAD4 inhibition, tumor cells were incubated in media from control or PAD4i-treated neutrophils that had been stimulated with CAF CMed. Media components from PAD4-inhibited neutrophils significantly impaired tumor cell growth compared to control conditions (Figure 3J). Together these data support the concept that t-NETs are pro-tumorigenic, and their inhibition is sufficient to prevent growth of established tumors through the restoration of an inflammatory state in tumor-infiltrating neutrophils.

The success of PAD4 inhibition in melanoma was not recapitulated in pancreatic tumors, with no significant changes in tumor volume being observed after treatment (Supplementary Figure 5A and B). However, systemic effects were detected, with PAD4 inhibition reducing the levels of clotting factors in the plasma (Supplementary Figure 5C) as with melanoma. Similarly, no significant changes in the stromal landscape within the tumor were detected (Supplementary figure 5D and E). We suggest that the lack of impact on these tumors, which still contained t-NETs (Figure 1B), may be due to the lack of drug penetration into the tumor which is a well-known phenomenon in mouse and human pancreatic cancer ^39,40^. However, treatment was effective as we observed reduced systemic clotting factors after PAD4 inhibition.

### CAF-derived Amyloid β drives t-NET formation

As neutrophils within the tumor traverse CAFs before entering the main tumor bulk, we next assessed whether direct contact was required for NET generation, or whether it was entirely soluble-mediator dependent as we observed systemically. To evaluate the capacity of CAFs to induce NETs by direct cell-cell interaction vs. secreted factors, neutrophils were seeded onto pancreatic tumor-derived CAFs in the presence or absence of FB or CAF CMed. While CAFs in the presence of CAF CMed readily induced NETosis, CAFs with normal FB media could not (Figure 4A). This suggested that direct contact between CAFs and neutrophils was not sufficient to drive t-NETosis, and the effect is primarily mediated through factors secreted by the CAFs.

Therefore, we performed proteomic analysis of the pancreatic FB and CAF CMed to identify the potential t-NET driver(s). Mass spectrometry revealed a number of differentially secreted proteins (Figure 4D and Supplementary figure 6) between these two cell types. Of interest, Amyloid β A4 protein (APP), fibronectin and heat shock protein 90 were significantly upregulated in CAF CMed (Figure 4E), all of which have been implicated in NETosis in different disease contexts ^41–44^. Treatment of neutrophils with a fibronectin inhibitor had no effect on the ability of CAF CMed to induce NETs (Supplementary figure 7A) thus we examined APP in greater detail. Similar to studies implicating Amyloid β peptide driven NETosis in Alzheimer’s ^45^, APP mRNA was detected in pancreatic CAFs at higher levels than pancreatic stellate cells (Supplementary figure 7B) and this was maintained at the protein level with Amyloid β detected in CAF CMed (Figure 4F). Upon determining that recombinant Amyloid β but not Amyloid α was able to induce NETosis in treated neutrophils in a dose dependent manner (Supplementary figure 7C) we next assessed whether Amyloid β was present in NET-rich tumors. In pancreatic tumors, two distinct patterns of APP distribution were observed with regions displaying either diffuse or punctate staining that co-localized with CAFs and tumor cells (Figure 4G) consistent with Amyloid β aggregates observed in other diseases. These data suggest that CAFs are a source of Amyloid β in the tumor, a key driver of NETosis.

To confirm that it was Amyloid β within CAF CMed that was responsible for the observed effects on neutrophils we inhibited the β-secretases which regulate secretion of Amyloid β (BACE 1-2). BACE inhibition in pancreatic and lung CAFs abolished the ability of the CMed to induce NETs *in vitro* (Figure 4H) supporting the hypothesis that Amyloid β production by CAFs underlies their capacity to induce t-NETs. A trend towards reduced neutrophil death was also observed in the absence of Amyloid β (Supplementary Figure 8A) indicating that ROS-mediated, suicidal NETosis in the tumor context is potentially driven by a single CAF-derived factor.

To further support our observations, I/V infusion of CAF CMed derived from CAFs pre-treated with BACE inhibitor to suppress Amyloid β production, abolished the systemic effects measured with CAF CMed (Figure 4I and Supplementary Figure 8B), whilst the effect of CAF CMed *in vivo* was recapitulated by infusion of recombinant Amyloid β alone (Figure 4I). We next examined the effects of BACE inhibition *in vivo* in skin tumour bearing mice (Figure 4J) using the same treatment regime as Cl-amidine and GSK484. As with inhibitors of NETosis, disruption of Amyloid β secretion prevented further increases in tumor volume compared with vehicle controls (Figure 4J), implying that CAF-driven Amyloid β-mediated NETosis supports tumour development *in vivo*, and that perturbation of either the NET driver or NET process is sufficient to prevent growth. This was confirmed using the B16.F10 melanoma model which is both neutrophil and NET poor ^35^. Treatment with CAF CMed or recombinant Amyloid β increased tumor growth (Figure 4K), potentially coinciding with enhanced infiltration of NETting neutrophils (Supplementary Figure 8D).

We then considered how CAF-derived Amyloid β may exert its effects on neutrophils. In light of reports that CD11b may act as a receptor for Amyloid β ^46^, we blocked CD11b during CAF CMed treatment and found that it completely abolished NETosis (Figure 4L and Supplementary Figure 8C) without affecting other functions such as phagocytosis (data not shown). In contrast, TLR2 neutralization on CAF CMed treated neutrophils had no effect on their capacity to NET (Figure 4M and Supplementary Figure 8C). Thus, CD11b on the surface of neutrophils may indeed function as a receptor, with its blockage desensitizing cells to the effects of Amyloid β present in CAF CMed.

### NETs reciprocally activate CAFs

Whilst we observed that t-NETosis was entirely soluble mediator driven (Figure 4A), the fact that NETs observed in the primary tumor site were often restricted to CAF dense regions (Figure 1A) led us to ask if t-NETs had reciprocal, pro-tumor effects on the phenotype and function of CAFs in their proximity. Treatment of CAFs with NETs supported an enhanced proliferation of CAFs (Figure 5A), and induced features of activation *in vitro* (Figure 5 B-E).

**Fig. 5.**
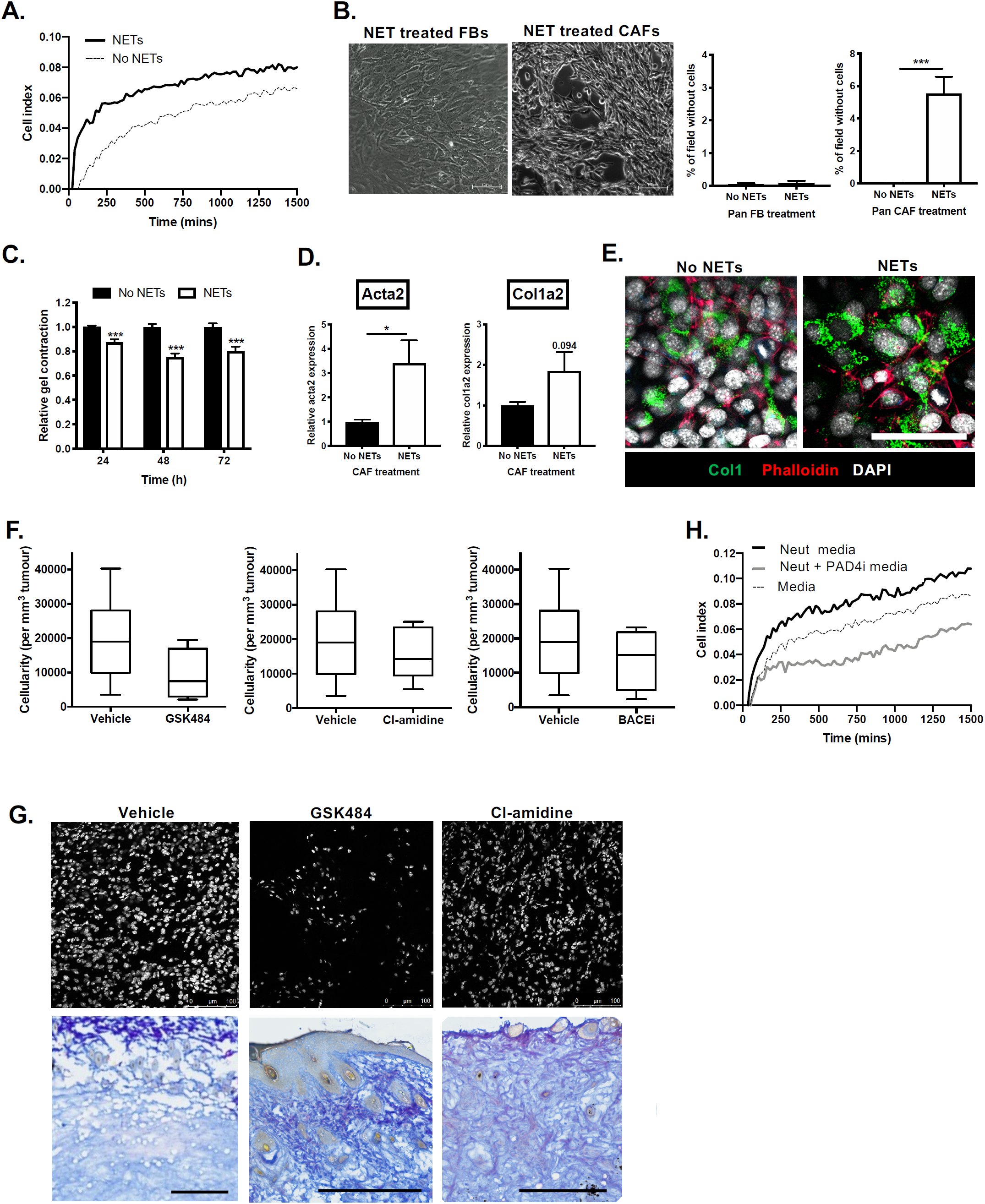
t-NETs induce CAF activation. (**A**) Representative plot illustrating growth of CAFs in the presence of vehicle or micrococcal nuclease detached NETs. (**B**) Phase contrast images and respective quantification of the percentage area of cell free space per field after treatment of pancreatic FBs or CAFs with NETs derived from CAF CMed treated neutrophils for 24h. (**C**) Quantification of the size of collagen gels 24h, 48h and 72h after seeding pancreatic CAFs treated with or without NETs derived from CAF CMed treated neutrophils for 24h. (**D**) Expression of Acta2 and Col1a2 at the gene level in pancreatic CAFs treated with or without NETs derived from CAF CMed treated neutrophils for 24h. (**E**) Confocal microscopy of Collagen1 and phalloidin on pancreatic CAFs treated with or without NETs derived from unstimulated neutrophils or CAF CMed treated neutrophils for 24h. (**F**) Cellularity of melanoma after treatment with GSK484, Cl-amidine, BACEi or vehicle expressed as cells per unit volume. (**G**) Representative confocal image of nuclei and collagen after treatment with GSK484, Cl-amidine, or vehicle. Tissues stained with Herovici stain to demark new collagen (blue) vs. mature collagen (pink). (**H**) Representative plot illustrating growth of CAFs following treatment with CAF CMed or CAF CMed and PAD4i-treated neutrophil-derived media. Data are mean ± SEM; * = p<0.05 and *** = p<0.001 using t-test. Assays were performed on (A and H) Representative of n=2 (in duplicate), (B) n=3 (in triplicate), (C) n=3 (in duplicate or quadruple), (D) n=4 (in duplicate) and (F) n=5-10 independent experiments. Scale bars are 50µm (B and D), 500µm (G upper panel) and 100µm (G upper panel).

We observed an increase in contraction of CAFs by the appearance of large gaps between CAFs treated with NETs derived from CAF CMed treated neutrophils compared to FB treated with NETs which remained as a monolayer (Figure 5B). To assess the contractile properties of the CAFs after treatment with t-NETs, we performed contraction assays. t-NET treated CAFs contracted collagen gels to a greater extent than CAFs treated with unstimulated neutrophil-derived NETs (Figure 5C), and coincided with an increased expression of αSMA and Col1a2 at the RNA and protein level (Figure 5D and E respectively). Examination of tumors treated with either PAD4 or BACE inhibitors revealed a reduced cellularity per unit volume when compared with vehicle treated tumours (Figure 5F and G). Moreover, consistent with earlier observations NET inhibition promoted degranulation and impaired tumor cell growth (Figure 3I and J), a similar effect was mirrored in CAFs (Figure 5H). Fibrotic structures with more mature collagen made up the remaining tumor mass (Figure 5G lower panel). Collectively, these data suggest that neutrophils skewed to undergo NETosis likely localise to CAF rich regions of a tumor where they form t-NETs in response to Amyloid β, promoting CAF expansion, contractility and deposition of matrix components supportive of tumor growth.

### Conservation of Amyloid β-NET axis in human disease

Having observed a significant pro-tumor communication between CAFs and neutrophils in multiple murine models, we sought to determine the clinical significance of these findings by examining human tumors. T-NETs were observed in human pancreatic tumors (Figure 6A), melanoma and melanoma metastases (Figure 6B and Supplementary Figure 9a), and when detected, both NETs and neutrophils were found juxtaposed to CAFs mirroring murine tumors. We next examined publicly available datasets for APP and β-secretase. Significantly, both pancreatic adenocarcinoma and cutaneous melanoma expressed elevated levels of APP and BACE2 compared to matched normal tissue (Figure 6C and D). When detected, elevated levels of Amyloid β could be measured circulating in the blood of patients with advanced melanoma compared with heathy controls (Supplementary Figure 9b). Moreover, expression of both app and bace2 strongly correlated with stromal markers pdpn, acta2, col1a2, cd34 typically used to identify CAFs, but not lymphatic marker lyve-1 (Figure 6E) inferring that correlations were specific to CAFs and not other components of the tumor stroma. These data recapitulate murine tumours, supporting CAFs as a source of Amyloid β in human pancreatic cancer and melanoma. The high expression of bace2, the rate-limiting step in Amyloid β release, rather than app was correlated with poorer patient prognosis in both tumor types (Figure 6F and G). Collectively, these findings suggest that Amyloid β secreted by CAFs is a critical driver of t-NET formation conserved in human cancers associated with poor prognosis, and importantly, has the potential to be detected in liquid biopsies.

**Fig. 6.**
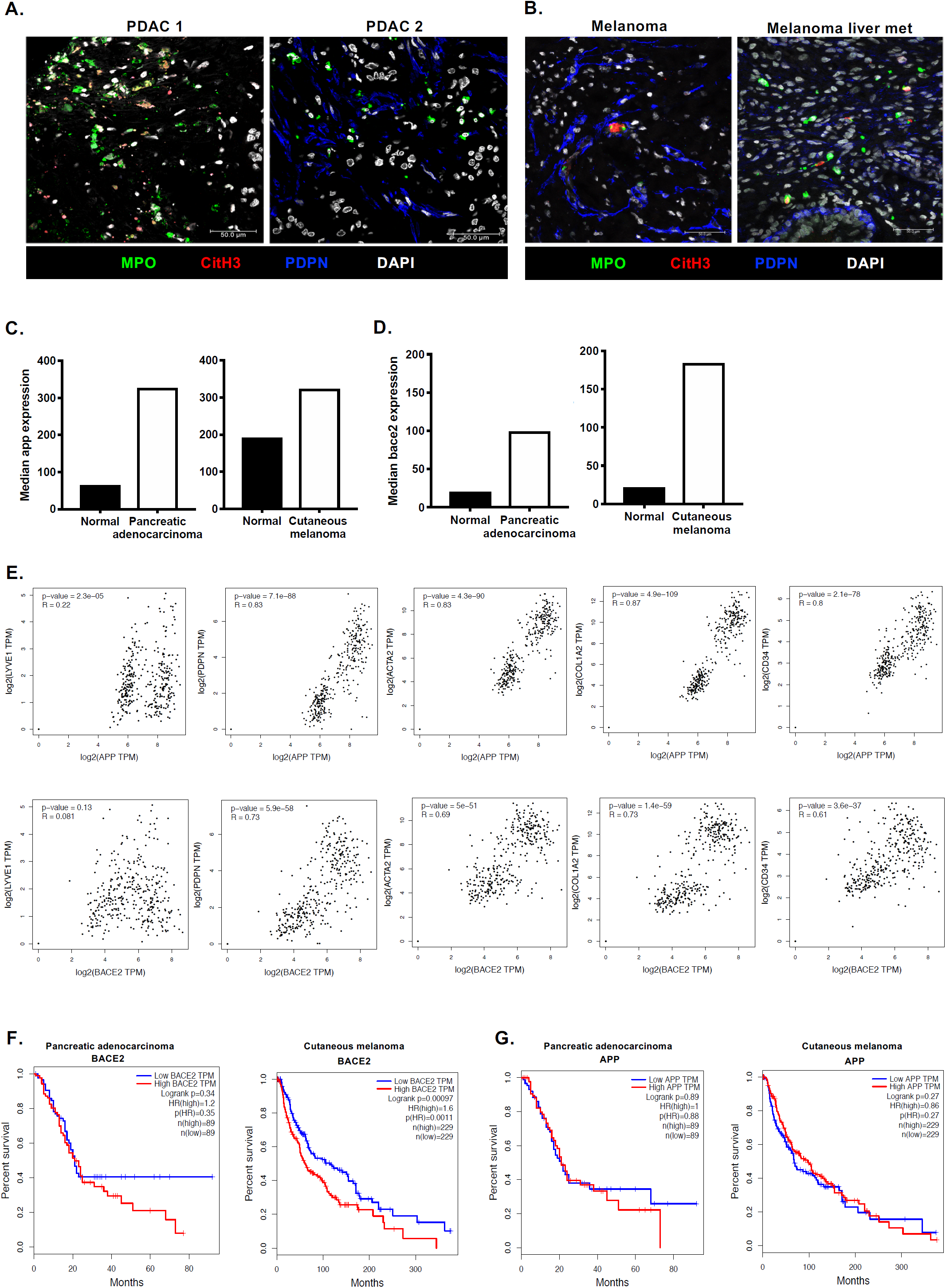
Conservation of t-NETs in human disease. (**A**) Representative confocal images of NETting neutrophils and CAFs in human pancreatic tumor biopsies (MPO, green; CitH3, red; podoplanin, blue; DAPI, white). (**B**) Representative confocal images of NETting neutrophils and CAFs in human melanoma primar tumor and metastasis samples (MPO, green; CitH3, red; podoplanin, blue; DAPI, white). (**C**) Median expression of app in human pancreatic adenocarcinoma and cutaneous melanoma compared to normal tissue from healthy donors and (**D**) Median expression of bace2 in human pancreatic adenocarcinoma and cutaneous melanoma compared to normal tissue from healthy donors *(GTEx and TCGA datasets)*. (**E**) Correlation of app and bace2 with a lymphatic marker (lyve1) and CAF markers (pdpn, acta2, col1a2 and cd34). Kaplin-meier curves showing overall survival of pancreatic adenocarcinoma and cutaneous melanoma patients correlated with high and low expression of **(F**) bace2 or (**G**) app.

## Discussion

The stroma is critical to tumour development and progression and it is evident that targeting stromal interactions offers many opportunities for cancer treatments ^1^. Here, we report the existence of a previously undescribed pro-tumor crosstalk between non-immune and immune stromal components to support growth. We demonstrate that neutrophils recruited to the tumor frequently localize to cancer-associated fibroblast (CAF)-rich areas, where they are stimulated to generate extracellular traps (NETs) supporting tumor growth. This process, which we have termed tumor-induced NETosis (t-NETosis), is driven by CAF-secreted Amyloid β both locally via CD11b on neutrophils within the primary tumor and at the systemic level within the bone marrow niche.

While neutrophil function has been extensively studied in the context of inflammation and tissue damage ^47–49^, there have been fewer studies relating to their contribution to the primary tumor ^32,50,51^ as well as the role of NETs and mediators responsible for driving their release ^21,51^. Contradictory studies have reported NET-derived proteins acting to both promote tumor cell proliferation and invasion ^52,53^, as well as inhibit tumor growth through apoptosis induction ^32,54^. In advanced disease, NETs have been reported to contribute to thrombus formation ^21^, metastatic colonization ^22,23^ and most recently the reactivation of dormant tumor cells ^29^. Previous studies point to a central role for G-CSF in recruiting neutrophils and to date is the only factor that has been identified as a NET inducer ^21,35^. This is particularly thought provoking given that cancer patients that are at high risk of becoming neutropenic receive G-CSF therapy in an attempt to restart myelopoiesis after chemo- and radiotherapy ^55^. Consistent with previous studies we found that CAF-induced NETosis was ROS and PAD4-dependent ^14^, but in this case did not require G-CSF.

With CAFs possessing such a potent effect on neutrophils via mechanisms distinct from previously described for systemic NETosis ^21^, we then examined the contribution of NETs in tumors. Inhibition of t-NETosis in mice with established tumors had no impact on neutrophil infiltration, yet was sufficient to abolish further growth. In these tumors, PAD4 inhibition translated to a skewing of neutrophils to a cytotoxic, pro-inflammatory and anti-tumor N1-like phenotype ^19,56^. Indeed, our data supported this hypothesis, confirming enhanced expression of activation markers and degranulation after PAD4 inhibition.

We subsequently identified Amyloid β, a protein typically associated with neurodegenerative disorders, as the critical CAF-derived factor inducing t-NETs. Inhibition of Amyloid β or its proposed cognate receptor, CD11b ^46^, prevented CAF-mediated NETosis and reduced growth of established tumors. Moreover, addition of soluble Amyloid β exacerbated tumor growth further implicating NETosis as pathological response in the cancer context. With the observed increase in expression of CD11b on neutrophils after CAF CMed treatment, our data imply that CAFs not only secrete the Amyloid β to drive NETosis, but may also render the neutrophils more responsive to circulating Amyloid β in tumor-bearing animals through increased expression of its receptor ^46^. This is in line with reports that Amyloid β can induce ICAM-1 expression by endothelial cells suggesting that it may play additional roles in the high affinity capture of infiltrating neutrophils ^41,57^.

In Alzheimer’s disease, Amyloid-driven NETosis has been associated with poor prognosis as a consequence of endothelial and parenchymal damage, and neurotoxicity ^57–59^. In this context, aggregation of Amyloid β not only drives the cognitive symptoms associated with Alzheimer’s, but its soluble form also acts as a DAMP contributing to the neuroinflammation which perpetuates progression ^60,61^. In cancer, we observed that Amyloid β secreted by CAFs form microaggregates in the tumor to drive NETosis *in situ* as well as disseminating into the blood where it conditions circulating and bone marrow resident neutrophils which raises the possibility that microaggregates of CAF-derived Amyloid β also act as a DAMP inducing aberrant NETosis to support tumor progression.

We provide evidence that neutrophils in pancreatic, skin and lung cancers exposed to CAF-derived cues exert pro-tumor effects that operate on multiple levels. Suppression of t-NETs in tumor bearing mice by inhibition of PAD4, or release of Amyloid β by BACE inhibition not only suppressed tumor growth but also brought about a decrease in thrombus formation as quantified by clotting factors; vWF and fibrinogen. With previous reports showing that NETs both contribute to thrombus formation, metastatic colonization and activation of dormant tumor cells ^21–23,29,35^ our findings that perturbation of either the driver or consequence of NETosis is sufficient to prevent tumor growth present the CAF-Amyloid-neutrophil axis as an attractive target. While the therapeutic potential of PAD4 blockade has yet to be clinically tested, it is possible that this approach may only provide a narrow therapeutic window since the inhibitor can acts when PAD4 is in a high calcium binding state ^62^, and may increase susceptibility to infection where NETosis is a critical response for bacterial clearance. Thus, inhibiting Amyloid β, for which drugs targeting BACE’s have already received FDA approval ^63^, and preventing neutrophils from receiving a NET stimulus may represent a more effective platform. Importantly, this would block pathological NETosis both at the primary tumor and systemically without impacting other critical neutrophil functions.

With the growing body of evidence supporting a role for NETs in advanced cancer - metastatic colonization and recurrence ^23,29,64^, and clinical data linking neutrophil infiltration with poor prognosis in multiple cancer types ^65–69^ – our findings warrant further studies to determine whether Amyloid β (circulating, or in the primary tumor) can be used as a biomarker to stratify patients. Although, in neuroinflammatory conditions such as Alzheimer’s disease, circulating Amyloid β is not currently considered to be a biomarker reflective of levels in the cerebrospinal fluid and central nervous system ^70^, we reported a trend towards an increase in Amyloid β levels in patients with melanoma compared to healthy volunteers. However, with our small cohort of archived patient samples, this was not significant. A number of external factors may be responsible for the observed variability; Circulating Amyloid β_1-40_ and β_1-42_ are predominantly bound to plasma proteins, and platelets or erythrocytes which can mask detection ^70^. Furthermore, differences in the hepatic and renal clearance of Amyloid β may influence the levels measured ^70^. Despite the difficulties in quantifying plasma Amyloid β levels, recent studies have implicated plasma Amyloid β in cancer. Higher levels of plasma Amyloid β were detected in hepatic cancer patients than those with Alzheimer’s disease ^71^, indicating that while cerebrospinal fluid levels are predictive of Alzheimer’s, circulating Amyloid β levels may be more reflective of cancer ^71^. Furthermore, it has been reported that breast and prostate tumor cells undergo amyloidosis as a result of environmental stress leading to cancer cell dormancy. This is particularly interesting in light of the recent study which implicated NETosis as a driver of dormant cancer cell activation in lung cancer ^72^. Indeed, amyloid accumulation in tumor cells may force the cells into a dormant state but as a consequence may also induce NETosis which could result in their activation and in a negative feedback system, re-initiation of the tumor. Amyloid β-induced NETosis drove tumor growth by reciprocally effecting CAFs, with t-NETs supporting proliferation and a more contractile phenotype. This phenomenon has also been reported in fibrosis where NETosis was found to drive activation, differentiation and the fibrotic response of human lung fibroblasts ^18^. Activation, matrix deposition and stiffening have all been correlated with disease progression and promotion of tumor growth in many cancers ^1,4^ without directly affecting tumor cell proliferation. Therefore, the direct action of Amyloid β on tumor cells previously reported ^72^ is distinct to the indirect effects it has on the tumor through the induction of NETosis further supporting a potential rationale for using BACE inhibitors to treat cancer.

In summary, we have demonstrated the existence of a novel mechanism by which CAFs stimulate t-NETosis at local and systemic levels via production of Amyloid β. Targeting this process with existing small molecule inhibitors stops tumour growth and restores a pro-inflammatory neutrophil phenotype. This sets the stage for further studies examining the potential of therapeutically targeting NETs.

## Methods

### Mice

All experiments involving animals were performed in accordance with UK Home Office regulations (PPL P88378375). For spontaneous genetic mouse tumors, Tyr::CreER; Braf^CA^; Pten^lox/lox^ (skin tumor) model, the LSL-KrasG12D/+;LSLTp53R172H/+;Pdx-1-Cre (pancreatic tumor) model, and inducible LSLKrasG12D/+;p53LSL-R270H/ER (lung tumor) model systems were tested. Orthotopic syngeneic tumours using B16.F10 melanoma were performed in approximately 8-week-old female C57BL/6 mice.

### PAD4 inhibition in vivo

Male and female mice were treated with PAD4 inhibitors; 3.5mM Cl-amidine (EMD Millipore), 20mg/kg GSK484 (Cayman Chemicals) or a vehicle control (DMSO). Where possible, technicians performing the experiment were blinded to drug treatments. Mice were recruited when tumors reached between 3-6mm in diameter and then received I/P doses of PAD4 inhibitor or vehicle control every day for 7d or until tumors reached their size limit. The size of pancreatic tumors was monitored by high-resolution ultrasound, as previously described ^2^. For skin tumors, the tumor volume was recorded daily using the formula (π/6)(shortest length*longest length)^2^. After treatment, the cellularity of the tumours was determined by flow cytometric analysis of the number of cells present in the tumour after digestion of the tissue. The number of cells was then normalised to tumor size. Plasma was isolated from skin and pancreatic tumor-bearing mice by collecting the blood from cardiac puncture and centrifuging at 800g for 10min. Plasma was then snap frozen for measurements of analytes at a later stage.

### BACE inhibition in vivo

Tumour-bearing Tyr::CreER; Braf^CA^; Pten^lox/lox^ mice were treated with 5mg/kg BACE inhibitor (Z-VLL-CHO, Abcam) or vehicle control. Mice were recruited when tumors reached between 3-6mm in diameter and then received I/P doses of BACE inhibitor for 7d or until tumors reached their size limit. Tumor volume was recorded daily using the formula (π/6)(shortest length*longest length)^2^. After treatment, the cellularity of the tumours was determined by flow cytometric analysis of the number of cells present in the tumour after digestion of the tissue. The number of cells was then normalised to tumor size.

### Amyloid β and CAF CMed treatment of tumors

2.5 × 10^5^ B16.F10 melanoma cells were inoculated subcutaneously into the shoulder region. Where possible, technicians performing the experiment were blinded to drug treatments; At day 5, day 7 and day 9 mice received I/P infusion of either vehicle, CAF CMed or recombinant Amyloid β. The tumor volume was recorded daily using the formula (π/6)(shortest length*longest length)^2^. After 11 days or when the tumors reached the size limit, the mice were sacrificed. The immune landscape and CAF composition of the tumors was analysed by flow cytometry.

### CAF CMed treatment in vivo

For investigations of CAF factors *in vivo* in the absence of tumors, CAF or FB CMed diluted 1:1 in complete culture media or 250μg/ml recombinant amyloid β was I/V infused into C57BL/6 mice and the bone marrow was harvested 24h later.

### Cell isolation and culture

CAFs were isolated from skin, lung *(58)* and pancreatic tumor bearing mice by mechanical separation of the tumor followed by digestion with an enzymatic cocktail consisting of 1mg/ml collagenase A and collagenase D and 0.4mg/ml DNase I (all from Roche) in PBS at 37°C for 2-3h with rotation at 600rpm. 10mM EDTA was then added to stop the enzymatic reaction. Normal FBs were isolated from matched tissues of wild type mice using the same method. CAFs and normal FBs were maintained in RPMI (Sigma-Aldrich) with 1.5 g/L NaHCO3, 10% fetal bovine serum (Life Technologies), 1% penicillin-streptomycin (Sigma-Aldrich), 10mM HEPES (Gibco), 15μM β-mercaptoethanol (Sigma-Aldrich).

### Conditioned media generation and fractionation

CMed from FB and CAFs was generated by culturing the cells until they reached 40-50% confluence and then changing the media to complete endothelial cell culture medium (Generon). The media was then harvested after 24h and filtered through 0.2um cell strainers before freezing. In some experiments, pancreatic or lung CAFs were treated with an inhibitor to beta-site Amyloid precursor protein cleaving enzyme 1 and 2 (BACE) (0.7μM; Abcam) for the duration of CAF CMed generation to inhibit Amyloid β production by the cells.

CAF CMed was also separated into the metabolite (less than 3kDa) and protein (more than 3kDa) fractions using 3kDa centrifugal filters (Merck Millipore) according to the manufacturer’s instructions. To isolate extracellular vesicles (MVs), CAF CMed was ultracentrifuged at 100,000g for 90min. MV depleted CMed was collected and the isolated MVs were resuspended in complete culture media.

### Neutrophil isolation

Wild type male and female C57BL/6 mice were sacrificed by cervical dislocation. Femurs and tibias were removed and BM aspirate was collected. Neutrophils were isolated using a two-step histopaque density gradient as previously described ^73^. Purified neutrophils were washed in PBS. To test the NETting capability of neutrophils from tumor bearing mice, bone marrow from mice bearing skin, lung and pancreatic tumors of varying size were isolated.

### NETosis assay

Bone marrow-derived neutrophils were counted and seeded onto poly-D-lysine coated plates at a density of 1×10^5^ cells per condition in serum free media. The media was changed to complete endothelial cell culture medium (Generon). To study NETosis, neutrophils were either treated with 1ug/ml Phorbol 12-myristate 13-acetate (PMA), CMed derived from FB or CAFs mixed 1:1 with complete culture media. Neutrophils were treated for 3h at 37°C and 5% CO_2_ and then stained with 20uM SYTOX(tm) Green Nucleic Acid Stain for 10min (Thermofisher Scientific). Images and videos were taken using a Zeiss Axio Observer.Z1 coupled with incubation chamber (ZEISS). 5 images were taken per well and each condition was performed in duplicate or triplicate.

To test the effects of ROS on NETosis, neutrophils were treated with anti-oxidants; 10mM n-acetyl cysteine (NAC), 20μM Diphenyleneiodonium (DPI), 2mM Trolox and 2mM Vitamin C for 30min prior to inducing NETosis and for the duration of the NET assay. Alternatively, neutrophils were treated with 0.03μg/ml α-G-CSF (R&D Systems) or 50μM Chloroquine (Sigma-Aldrich) for the duration of the NET assay. In some experimemts, neutrophils were treated with 1.5mg/ml Cl-amidine, an inhibitor of the Protein Arginine Deiminase 4 (PAD4), 10μg/ml anti-CD11b or anti-TLR2 (Biolegend) or 1μg/ml fibronectin inhibitor (Santa-Cruz Biotechnology) for the duration of culture. The area of NET coverage was quantified using ZEN Lite (ZEISS) and ImageJ (Fiji) software by drawing around every NET within a field. Dead cells (positive for SYTOX green) were also counted per field.

### NET isolation and quantification

After NET generation, neutrophils were treated with 1U/ml micrococcal nuclease (Sigma-Aldrich) for 10min (found to be the optimum time for detaching NETs from neutrophils without digesting the NETting DNA) at 37°C and 5% CO_2_. The enzyme was inactivated with 0.5mM EDTA.

### Gel contraction assays

Pancreatic CAFs were treated with approximately 10ng/ml NETting DNA (taken from 1×10^5^ neutrophils stimulated with CAF CMed or PMA) for 24h or from untreated neutrophils. CAFs were trypsinized and then seeded into 2mg/ml collagen gels (Rat tail collagen, BD Biosciences) at a density of 1×10^5^ cells/gel in 24 well plates. Gels were left to polymerise for 20min at 37 °C before adding full media. The gel was detached from the culture dish using a pipette tip. The gels were imaged at 24h, 48h and 72h after generation. The area of the gel was then measured using ImageJ software. The relative gel area was then calculated by comparing it to gels containing untreated CAFs.

### Flow cytometry

#### Tumor digestion

Tumors were minced using a razor and digested with 1mg/ml collagenase A and collagenase D and 0.4mg/ml DNase I in PBS at 37°C for 2h with rotation at 600rpm. 10mM EDTA was then added to stop the enzymatic reaction. The cell suspension was passed through a 70μm filter and stained with live/dead fixable violet stain (Thermofisher Scientific). Cells were subsequently stained with the following fluorescently conjugated antibodies; CD45 (30-F11), Ly6G (1A8), F4/80 (BM8), CD11b (M/170), CD11c (N418), Thy1 (30-H12), Podoplanin (8.1.1.), PDGFRα (APA5; all from Biolegend) and CD31 (390; eBioscience) at 1:300 dilution. Flow cytometry was performed on LSR Fortessa (BD Biosciences) analyzers. Unstained and single-stained compensation beads (Invitrogen) were run alongside to serve as controls. Offline analysis was carried out on FlowJo (Treestar). For *in vivo* PAD4 inhibitor studies, tumors were separated for flow cytometric and immunofluorescent analysis. Some GSK484 treated tumors were too small for analysis and were therefore excluded (only tumor volumes were recorded).

### In vitro treatment of skin tumor cells with PAD4 inhibitors

Tumor cells were isolated from skin tumors by digestion with 4mg/ml Collagenase A for 1h at 37°C. The cells were then strained through a filter and seeded onto a culture dish in DMEM supplemented with 5% FBS and 1% penicillin/streptomycin. The cells were cultivated for 3-4d and then seeded onto 24-well plates at a density of 3×10^4^ cells/well. The cells were either treated with DMSO (vehicle control), 100µM Cl-amidine or 10µM GSK484 for 48h. Images were taken at 0h, 24h and 48h after treatment.

### In vitro neutrophil staining

Bone marrow neutrophils were isolated as described above and treated with lung FB or CAF CMed for 30min. Neutrophils were then treated with 10μM 2′,7′-Dichlorodihydrofluorescein diacetate (DCFDA; Sigma-Aldrich) or 2×10^8^ yellow/green 1μm fluoresbrite beads (Polysciences Inc.) for 20min. Alternatively, neutrophils were treated with CMed and stained with antibodies for CD11b (M/170), CD18 (M18/2) and CD62L (MEL-14; all from Biolegend). In some experiments, neutrophils were treated with CMed and stained with Annexin V (BD Pharmingen) and 7-AAD (Thermofisher Scientific) for 30min. For all experiments, the cells were washed and then immediately analyzed by flow cytometry.

### PAD4 inhibitor treatment on neutrophil function in vitro

Murine bone marrow neutrophils were seeded at 1×10^5^ cells per condition in complete endothelial cell culture medium mixed 1:1 with pancreatic CAF CMed. Neutrophils were treated with or without 1.5mg/ml Cl-amidine for 3h. Neutrophil activation was then assessed by measuring expression of CD11b, CD18 and CD62L. ROS production was assessed by measuring DCFDA by flow cytometry (as above). The phagocytic activity of the neutrophils was assessed by measuring fluorescent bead uptake by flow cytometry. Neutrophil degranulation was determined by surface expression of CD35 and CD63.

### Characterization of CAFs and FB

Isolated CAFs and FB were stained for typical markers; Podoplanin, PDGFRα and Thy1 (as described above) and markers to exclude immune cells (CD45), endothelial cells (CD31) and epithelial cells (EpCAM clone G8.8; all from Biolegend).

### Gene expression analysis

Pancreatic CAFs were treated with approximately 10ng/ml NETting DNA (from CAF CMed or PMA stimulated neutrophils) for 24h. RNA was isolated using the RNeasy Mini Kit (Qiagen, Crawley, UK), converted to cDNA and analyzed by qPCR using Universal PCR mastermix (Life Technologies) according to manufacturer’s instructions. Primers were bought as Assay on Demand kits from Applied Biosystems. qRT-PCR was performed using TaqMan assays (*Col1a2* Mm00483888_m1, *Acta2* Mm00725412_s1 and *Gapdh* Mm99999915_g1) and a StepOne Real Time PCR System (both Life Technologies). Levels of each gene was expressed as 2^-δCT^ (relative to Gapdh).

### Analysis of Amyloid-related genes in human tumors

Expression of app and bace2 and correlations with CAF and lymphatic markers in pancreatic adenocarcinoma and skin cutaneous melanoma compared to healthy tissue were analyzed using GEPIA RNA-sequencing expression data taken from the TCGA and GTEx projects ^74^.

### Immunofluorescence

Tumors were snap frozen in OCT medium (TissueTek). 5-7µm sections were fixed in ice cold acetone/methanol for 5min. Sections were blocked in 10% donkey serum in PBS and incubated with primary antibodies overnight at 4°C. Primary antibodies as follows: Rabbit anti-citrullinated histone H3 R17+R2+R8 (ab5103, 1:500, Abcam), goat anti-myeloperoxidase (AF3667, 1:1000, RnD Systems), hamster anti-podoplanin (clone 8.1.1, 127402, 1:100, BioLegend), Alexa 488 conjugated mouse anti-APP (clone 22C11, MAB348A4, 1:100, Merck-Millipore), mouse anti-αSMA (clone 1A4, MAB1420, 1:100, RnD Systems). Slides were washed, incubated with appropriate secondary antibodies and counterstained with DAPI. Sections were mounted in Slowfade Gold (Invitrogen). Images were acquired on a Leica SP5 confocal microscope.

Paraffin-embedded formalin fixed human pancreatic TMA slides (Biomax) (LREC HBREC 2019.16) were dewaxed in xylene and rehydrated in graded alcohols prior to antigen retrieval in Tris-EDTA pH9. Slides were blocked and incubated with primary antibodies as above at 4°C overnight. Samples were washed and incubated in fluorescently conjugated secondary antibodies before cell nuclei were counterstained with DAPI and sections were mounted in Slowfade Gold and imaged on the Leica SP5.

### ELISA

Levels of murine Amyloid β42, (Fisher Scientific), von Willebrand factor (vWF) and fibrinogen (both from Abcam) were measured in plasma collected from skin and pancreatic tumor bearing mice or in serum-free CMed from pancreatic CAFs as per manufacturer’s instructions. Human platelet poor plasma was obtained by centrifuging whole blood from healthy donors at 2000g for 15min. Samples were immediately snap frozen until use. Archived plasma samples were obtained from melanoma patients (MELRESIST NRES: 11.NE.0312). Levels of Amyloid β42, von Willebrand factor (vWF) and fibrinogen (all from Abcam) in human melanoma patients and healthy controls was assessed as per manufacturers guidelines.

### Mass spectrometry

FB and CAF CMed were generated without fetal bovine serum for 24h. The protein fractions were isolated and concentrated using 3kDa centrifugal filters (Merck Millipore). LC MS/MS was performed on the concentrated culture CMed and spectral analysis was performed. Data was analyzed using Scaffold 4 software.

### Measurement of NETs or NET CM on cells in vitro

#### NETs

1×10^5^ neutrophils were treated with CAF CMed diluted 1:1 in complete EC media (Cell Biologics) with or without Cl-amidine and incubated for 3h to generate NETs. The media was then harvested from the NETting or NET inhibited neutrophils. PBS supplemented with 1U/ml micrococcal nuclease (Sigma-Aldrich) was then added to the neutrophils for 10min at 37°C and 5% CO_2_ to detach the NETs. The enzyme was inactivated with 0.5mM EDTA. PBS containing the NETs was then harvested for xCELLigence assays.

#### Neutrophil-derived factors

As for above, media from NETting or NET inhibited neutrophils was harvested for xCELLigence assays. Pancreatic CAF or tumour cells were seeded onto 16 well E-plates (ACEA Biologics) at a density of 5×10^3^ cells/well. Cells were allowed to adhere for 1h. The media was then replaced with the NETting or NET inhibited neutrophil conditioned media or the corresponding NETs. Plates were placed into an xCELLigence RTCA MP Real Time Cell Analyzer (ACEA Biologics). Recordings of the impedance, correlating to cell proliferation, were then taken over a 48h period. The background was then subtracted from the appropriate wells.

### Statistical analysis

Data are expressed as mean ± SEM, where a different neutrophil isolate and batch of CMed was used for each experiment. Multi-variant data were analyzed using analysis of variance (ANOVA), followed by Dunnett or Tukey post-hoc tests. Mann-Whitney or t-test was used to compare individual treatment conditions. p<0.05 was considered statistically significant. For mass spectrometry, differences in protein expression in FB and CAF CMed was considered significant when the spectral count was p<0.01.

## Acknowledgments

The authors would like to thank staff at the ARES and CRUK Cambridge Institute animal facility for assistance with *in vivo* experiments, and members of the CIMR flow cytometry core for assistance with flow cytometry applications. This work was supported by Medical Research Council Core funding. T.J. was supported by Cancer Research UK funding, Clinician Scientist grant (C42738/A24868); Cold Spring Harbor Laboratory (CSHL) and Northwell Health for unrestricted funding, and US National Institute of Health for funding received as part of Cancer Center Support Development Funds granted to CSHL (5P30CA045508-31)

## Author contributions

J.D.S. and H.M. conceived the study, designed, performed, analysed and interpreted experiments, and wrote the manuscript. J.J performed experiments and critically edited the manuscript. T.J. performed KPC experiments and provided advice and critically edited the manuscript

## Data and materials availability

Publicly available data were obtained from TCGA and the GTEx projects and the GEO repository (accession number GSE42605). All data is available in the main text or the supplementary materials upon reasonable request.

