## Supplementary figures and legends for "Stromal Amyloid β drives Neutrophil extracellular trap formation to augment tumour growth"

**Supplementary Figure Legends**

**Fig. S1.** Neutrophil activation and function are unaltered by CAF-derived factors. (**A**) Characterization of skin, lung and pancreatic FBs and CAFs by flow cytometry showing that the cells express the classic CAF markers (Thy1, Podoplanin, PDGFR$\alpha$) on the surface and lack expression of tumor cell (EpCAM) and immune cell (CD45) markers. (**B**) Quantification of neutrophil death after treatment with FB CMed, CAF CMed from lung or pancreas or PMA based on expression of Annexin V and 7-AAD by flow cytometry. (**C**) Quantification of CD11b, CD18 and CD62L expression on neutrophils after treatment with lung FB or CAF CMed with and without simulation with LPS by flow cytometry. (**D**) Quantification of ROS production by neutrophils after treatment with lung CAF CMed or PMA based on the levels of DCFDA by flow cytometry. (**E**) The phagocytic capacity of neutrophils after lung CAF CMed or PMA treatment based on uptake of fluorescent 1µm beads assessed by flow cytometry. Data are mean ± SEM; * = p<0.05 and ** = p<0.01 using t-test. Assays were performed on (A) n=1, (B) n=2, (C) n=4, (D) n=5-8 and (E) n=5 independent experiments.

**Fig. S2.** CAF induced NETosis is a ROS-dependent, autophagy independent response. (**A**) Schematic of NETosis assay (**B**) Representative phase contrast images of SYTOX green stained dead neutrophils and NETs after 3h treatment with PMA or pancreatic CAF CMed. Inset; zoomed image of a NET (green haze) outlined with a dashed white line. Quantification of the relative NET coverage of neutrophils treated with lung CAF CMed with or without treatment with (C) Chloroquine (Cl), (D) Diphenyleneiodonium (DPI), (E) Trolox and (F) Vitamin C. Data are mean ± SEM; * = p<0.05, ** = p<0.01 and *** = p<0.001 using t-test. Assays were performed on (C) n=5, (D) n=4, (E-F) n=2 (in triplicate) independent experiments. Scale bars correspond to 50µm.

**Fig. S3.** CAF-derived factors drive NETosis systemically. (A) Quantification of the percentage death of neutrophils of bone marrow neutrophils taken from mice with pancreatic, lung or skin tumors. (B) Quantification of percentage death of bone marrow neutrophils taken from wild type mice intravenously infused with pancreatic or lung FB or CAF CMed 24h before performing the NETosis assay. (C) Quantification of percentage death and relative number of bone marrow neutrophils taken from mice intravenously infused with lung CAF CMed with or without pre-treatment with Cl-amidine for 24h. Data are mean ± SEM; * = p<0.05, ** = p<0.01 and *** = p<0.001 using (A and C) t-test and (B) one-way ANOVA with Dunnett post hoc test. Assays were performed on (A) n=4 (in duplicate), n=7 (in quadruplet) and n=7 for pancreatic, lung and skin tumor bearing mice respectively, (B) n=8 (in triplicate) and n=6-16 (in duplicate) for pancreatic and lung FB and CAF CMed infused mice respectively, (C) n=8 (in triplicate) independent experiments.

**Fig. S4**. Treatment of skin tumor bearing mice with a broad-spectrum PAD inhibitor reduces tumor growth. (A) Relative volume of skin tumors on mice treated with vehicle or Cl-amidine over 7d. (B) Flow cytometric analysis of the percentage immune cells ($CD45^+$), myeloid cells ($CD11b^+$), macrophages ($F4/80^+$) and neutrophils ($Ly6G^+$) in the tumors treated with Cl-amidine. (C) Flow cytometric analysis of CAF populations in the tumors treated with Cl-amidine based on the percentage Thy1, Podoplanin and PDGFRα positive cells. (D) The levels of clotting factors (fibrinogen and von Willerbrand factor; vWF) in the plasma of skin tumor bearing mice treated with vehicle or Cl-amidine. (**E**) Flow cytometric analysis of CAF populations in the tumors treated with GSK484 based on the percentage Thy1, Podoplanin and PDGFRα positive cells. (F) Representative phase contrast images of skin tumor cells after treatment with vehicle, 100µM Cl-amidine or 10µM GSK484 over 48h. Data are mean ± SEM; * = p<0.05 and ** = p<0.01 using a Mann-Whitney test. Assays were performed on (A-D) n=7-8, (E) n=4-6, (F) n=6-7 independent experiments.

**Fig. S5.** Treatment of pancreatic tumor bearing mice with a broad-spectrum PAD inhibitor has no effect on tumor growth. Schematic of Cl-amidine treatment regime for pancreatic tumor bearing mice. (B) Relative cross-sectional area (CSA) and diameter of tumors treated with vehicle or Cl-amidine over 7d. (C) The levels of clotting factors (fibrinogen and von Willerbrand factor; vWF) in the plasma of pancreatic tumor bearing mice treated with vehicle or Cl-amidine. (D) Flow cytometric analysis of the percentage immune cells ($CD45^+$), myeloid cells ($CD11b^+$), macrophages ($F4/80^+$) and neutrophils ($Ly6G^+$) in the tumor. (E) Flow cytometric analysis of CAF populations in the tumor based on the percentage Thy1, Podoplanin and PDGFRα positive cells. Assays were performed on n=4 independent experiments.

**Fig. S6.** Enlarged heatmap for differentially secreted proteins in pancreatic FB and CAF CMed analyzed by mass spectrometry

**Fig. S7.** Amyloid β is sufficient to drive NETosis *in vitro*. (**A**) Quantification of the relative NET coverage and percentage death of neutrophils treated with pancreatic CAF CMed with or without a fibronectin inhibitor (FNi). (**B**) Log2(reads per kilobase transcript per million map reads; RPKM+1) expression of the app gene in pancreatic (pan) CAFs normalized to pancreatic (pan) stellate cells from publicly available data *(2)*. (**C**) Quantification of the relative NET coverage and percentage death of neutrophils treated with recombinant Amyloid α (Aα), recombinant Amyloid β (Aβ) or PMA for 3h. Data are mean ± SEM; * = $p<0.05$, ** = $p<0.01$ and *** = $p<0.001$ using (A) t-test and (C) one-way ANOVA with a Dunnett post hoc test. Assays were performed on (A) n=2 (in duplicate), (C) n=3 (in duplicate or triplicate) independent experiments.

**Fig. S8.** Inhibition of Amyloid β secretion by CAFs suppresses NETosis (A) Quantification of percentage death of neutrophils treated with pancreatic and lung CAF CMed generated with or without 24h pre-treatment of the CAFs with β-secretases 1 and 2 (BACE1-2) inhibitor.(B) Quantification of the percentage death of bone marrow neutrophils taken from wild type mice 24h after intravenous infusion with pancreatic CAF CMed taken from cells pre-treated with or without BACE1-2 inhibitor for 24h or recombinant Amyloid β. (C) Quantification of cell death after 3h treatment with pancreatic CAF CMed with or without CD11b or TLR2 blocking antibodies. (D) Representative confocal images showing neutrophils (myeloperoxidase (MPO; in green), CAFs (podoplanin, in blue) and counterstained with DAPI (in white) of orthotopically implanted B16.F10 tumor cells with vehicle, CAF CMed or recombinant Amyloid β treatment. Assays were performed on (A) n=6 (in triplicate), (B) n=7 (in duplicate or triplicate), (C) n=3 (in triplicate).

**Fig. S9.** Representative confocal images of NETting neutrophils and CAFs in a human brain metastasis (MPO, green; CitH3, red; podoplanin, blue; DAPI, white). (B) Quantification of Amyloid β in plasma from melanoma patients and healthy controls. Data are mean ± SEM. Assays were performed on (B) n=10-20 independent experiments.

**Movie S1.**

Neutrophil stained with SYTOX green undergoing NETosis.  Net release marked by a “sudden haze” of green material.

Figure 1

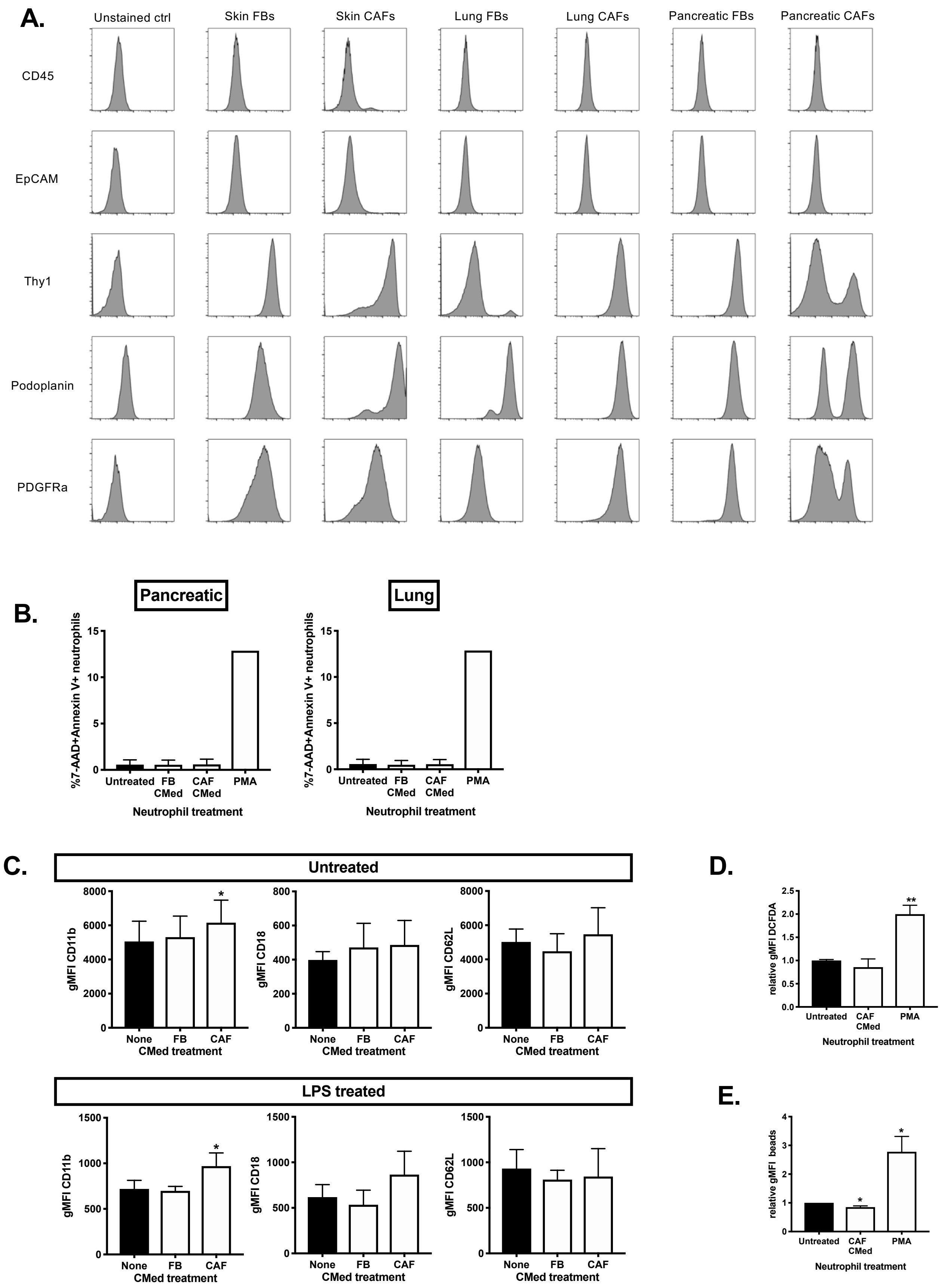

A.
Unstained ctrl
Skin FBs
Skin CAFs
Lung FBs
Lung CAFs
Pancreatic FBs
Pancreatic CAFs
CD45
EpCAM
Thy1
Podoplanin
PDGFRa
B.
Pancreatic
Lung
%7-AAD+Annexin V+ neutrophils
Untreated
FB CMed
CAF CMed
PMA
Neutrophil treatment
C.
Untreated
gMFI CD11b
gMFI CD18
gMFI CD62L
None
FB
CAF
CMed treatment
LPS treated
D.
relative gMFI DCFDA
Untreated
CAF CMed
PMA
Neutrophil treatment
E.
relative gMFI beads

### Figure S2

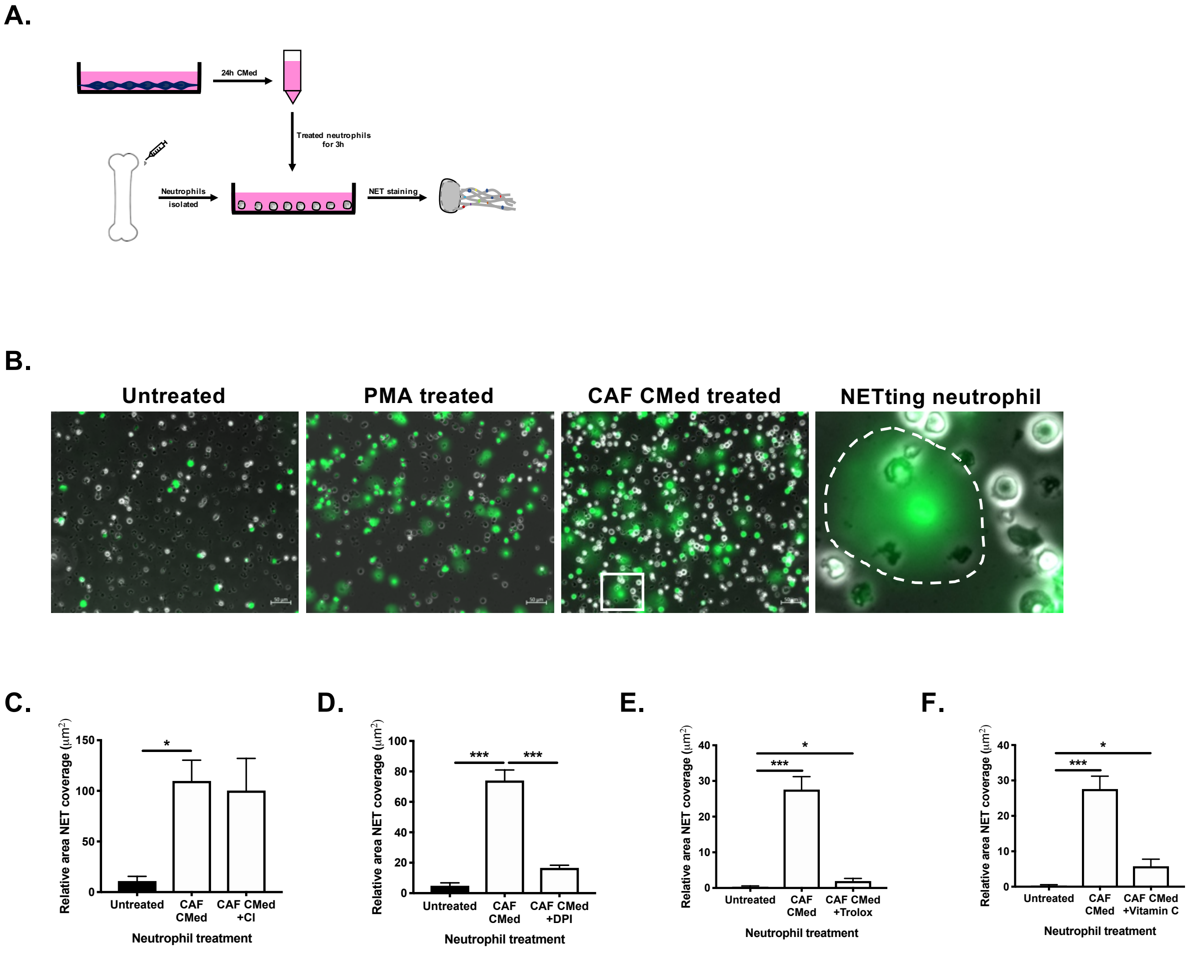

A.
24h CMed
Treated neutrophils for 3h
Neutrophils isolated
NET staining
B.
Untreated
PMA treated
CAF CMed treated
NETting neutrophil
C.
Relative area NET coverage (μm²)
150
100
50
0
*
Untreated
CAF CMed
CAF CMed +CI
Neutrophil treatment
D.
Relative area NET coverage (μm²)
100
80
60
40
20
0
***
***
Untreated
CAF CMed
CAF CMed +DPI
Neutrophil treatment
E.
Relative area NET coverage (μm²)
40
30
20
10
0
*
***
Untreated
CAF CMed
CAF CMed +Trolox
Neutrophil treatment
F.
Relative area NET coverage (μm²)
40
30
20
10
0
*
***
Untreated
CAF CMed
CAF CMed +Vitamin C
Neutrophil treatment

**Figure S3**

**A.**

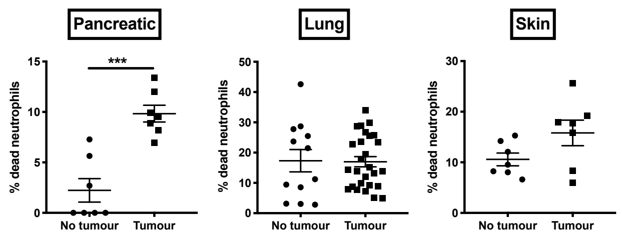


**B.**

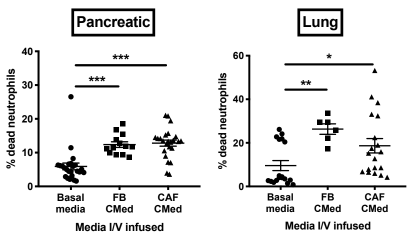


**C.**

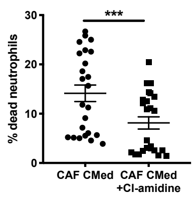

### Figure S4

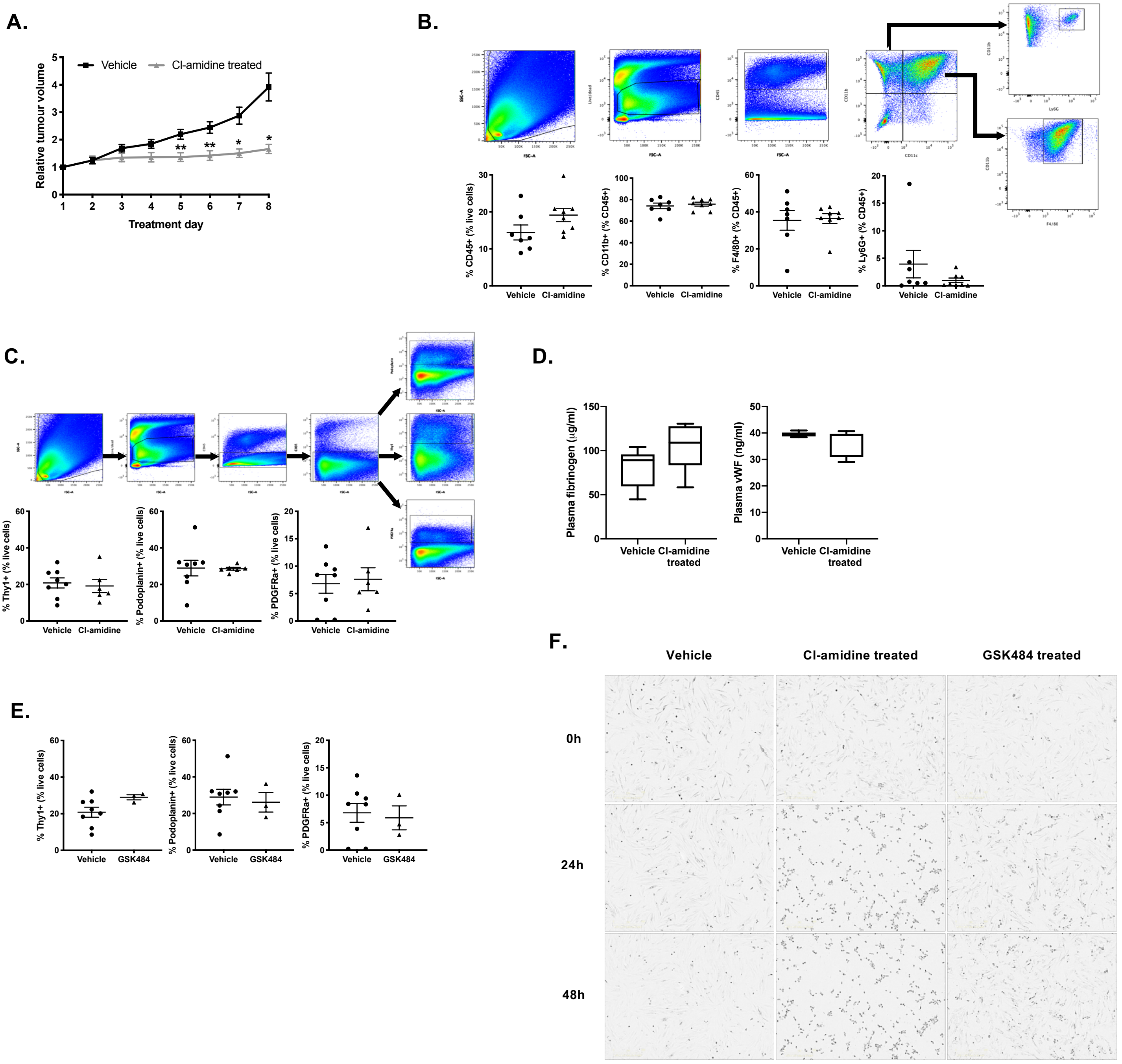

A.
Vehicle
Cl-amidine treated
Relative tumour volume
Treatment day
B.
% CD45+ (% live cells)
% CD11b+ (% CD45+)
% F4/80+ (% CD45+)
% Ly6G+ (% CD45+)
Vehicle
Cl-amidine
C.
% Thy1+ (% live cells)
% Podoplanin+ (% live cells)
% PDGFRa+ (% live cells)
Vehicle
Cl-amidine
D.
Plasma fibrinogen (μg/ml)
Plasma vWF (ng/ml)
Vehicle
Cl-amidine treated
E.
% Thy1+ (% live cells)
% Podoplanin+ (% live cells)
% PDGFRa+ (% live cells)
Vehicle
GSK484
F.
Vehicle
Cl-amidine treated
GSK484 treated
0h
24h
48h

#### Figure S4

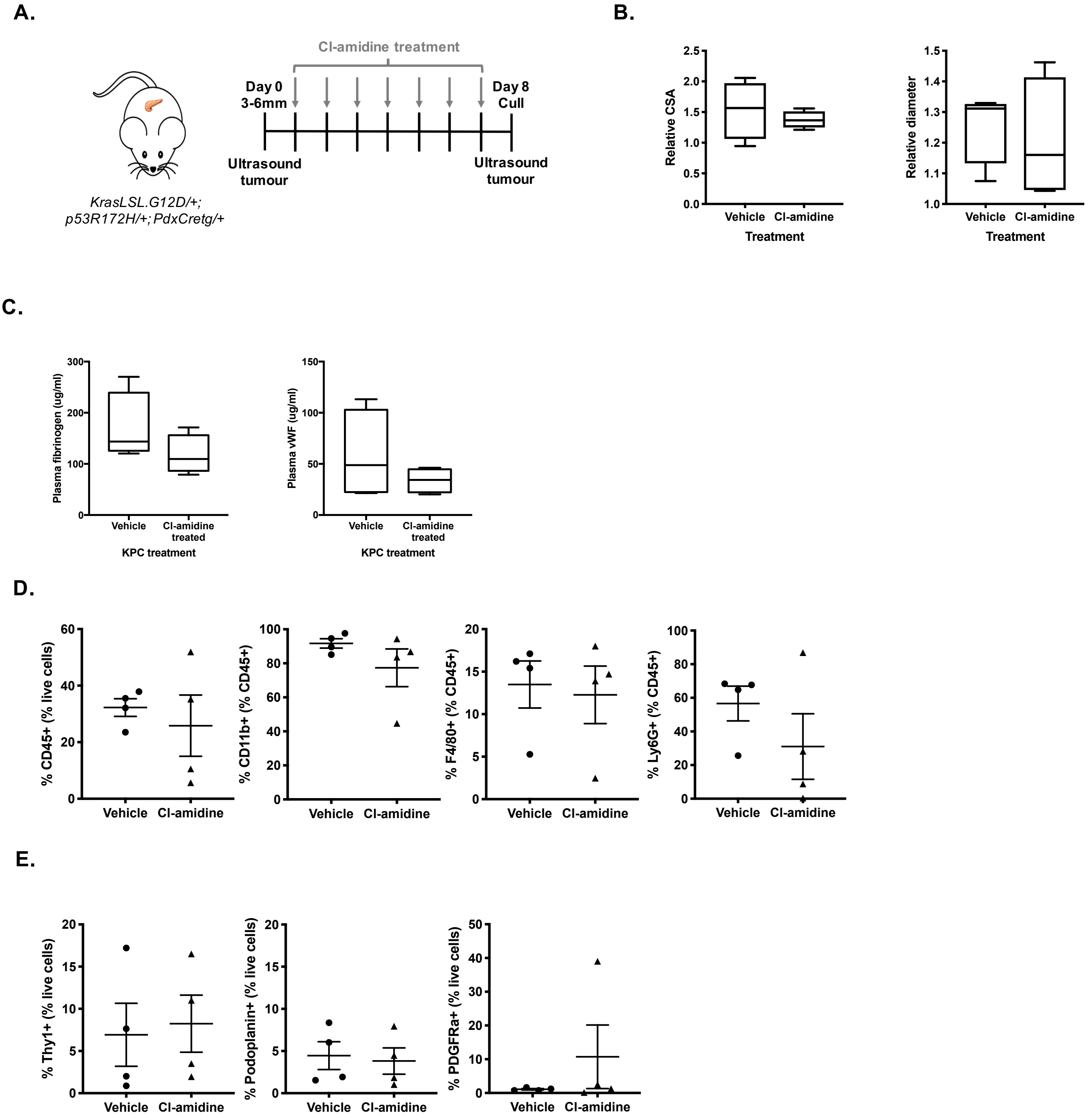
A.
Cl-amidine treatment
Day 0
3-6mm
Day 8
Cull
Ultrasound
tumour
Ultrasound
tumour
KrasLSL.G12D/+;
p53R172H/+; PdxCretg/+
B.
Relative CSA
Vehicle
Cl-amidine
Treatment
Relative diameter
Vehicle
Cl-amidine
Treatment
C.
Plasma fibrinogen (ug/ml)
Vehicle
Cl-amidine
treated
KPC treatment
Plasma vWF (ug/ml)
Vehicle
Cl-amidine
treated
KPC treatment
D.
% CD45+ (% live cells)
% CD11b+ (% CD45+)
% F4/80+ (% CD45+)
% Ly6G+ (% CD45+)
Vehicle
Cl-amidine
E.
% Thy1+ (% live cells)
% Podoplanin+ (% live cells)
% PDGFRa+ (% live cells)
Vehicle
Cl-amidine

### Figure S6

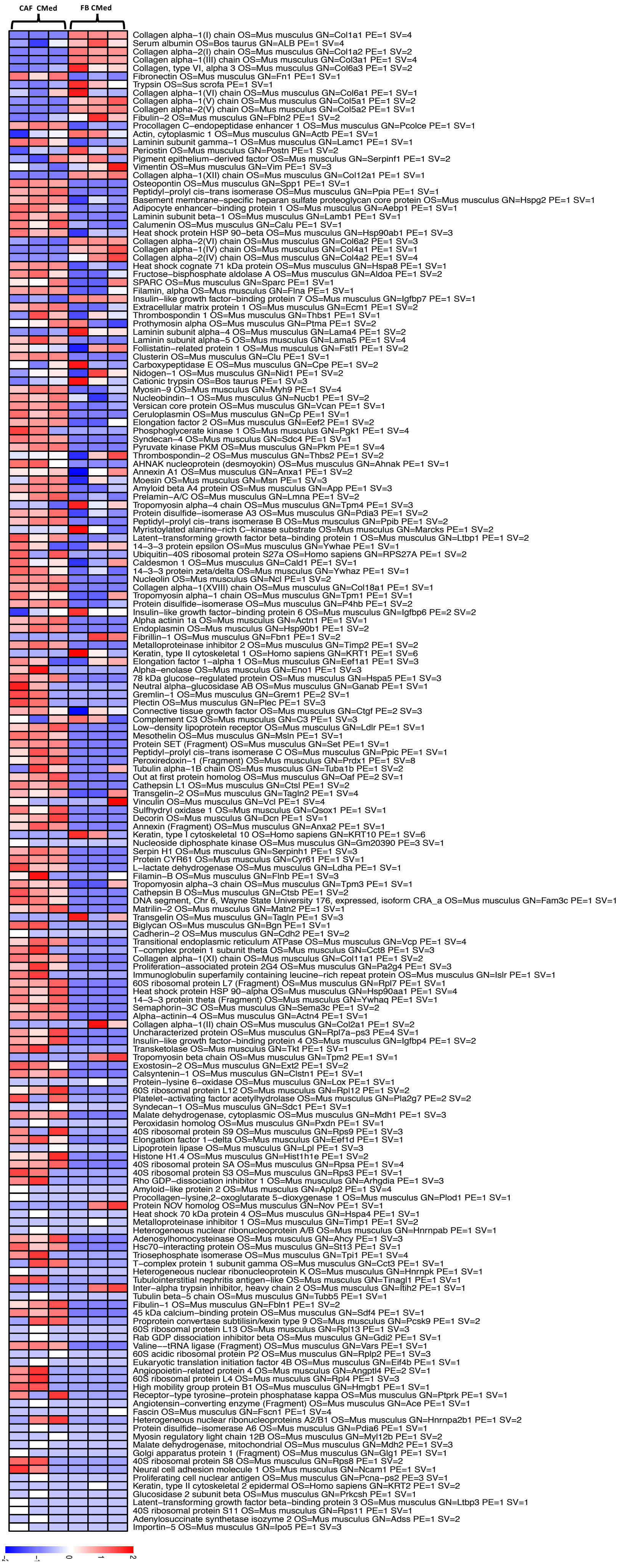
CAF CMed
FB CMed
Collagen alpha−1(I) chain OS=Mus musculus GN=Col1a1 PE=1 SV=4
Serum albumin OS=Bos taurus GN=ALB PE=1 SV=4
Collagen alpha−2(I) chain OS=Mus musculus GN=Col1a2 PE=1 SV=2
Collagen alpha−1(III) chain OS=Mus musculus GN=Col3a1 PE=1 SV=4
Collagen, type VI, alpha 3 OS=Mus musculus GN=Col6a3 PE=1 SV=2
Fibronectin OS=Mus musculus GN=Fn1 PE=1 SV=1
Trypsin OS=Sus scrofa PE=1 SV=1
Collagen alpha−1(VI) chain OS=Mus musculus GN=Col6a1 PE=1 SV=1
Collagen alpha−1(V) chain OS=Mus musculus GN=Col5a1 PE=1 SV=2
Collagen alpha−2(V) chain OS=Mus musculus GN=Col5a2 PE=1 SV=1
Fibulin−2 OS=Mus musculus GN=Fbln2 PE=1 SV=2
Procollagen C−endopeptidase enhancer 1 OS=Mus musculus GN=Pcolce PE=1 SV=1
Actin, cytoplasmic 1 OS=Mus musculus GN=Actb PE=1 SV=1
Laminin subunit gamma−1 OS=Mus musculus GN=Lamc1 PE=1 SV=1
Periostin OS=Mus musculus GN=Postn PE=1 SV=2
Pigment epithelium−derived factor OS=Mus musculus GN=Serpinf1 PE=1 SV=2
Vimentin OS=Mus musculus GN=Vim PE=1 SV=3
Collagen alpha−1(XII) chain OS=Mus musculus GN=Col12a1 PE=1 SV=1
Osteopontin OS=Mus musculus GN=Spp1 PE=1 SV=1
Peptidyl−prolyl cis−trans isomerase OS=Mus musculus GN=Ppia PE=1 SV=1
Basement membrane−specific heparan sulfate proteoglycan core protein OS=Mus musculus GN=Hspg2 PE=1 SV=1
Adipocyte enhancer−binding protein 1 OS=Mus musculus GN=Aebp1 PE=1 SV=1
Laminin subunit beta−1 OS=Mus musculus GN=Lamb1 PE=1 SV=1
Calumenin OS=Mus musculus GN=Calu PE=1 SV=1
Heat shock protein HSP 90−beta OS=Mus musculus GN=Hsp90ab1 PE=1 SV=3
Collagen alpha−2(VI) chain OS=Mus musculus GN=Col6a2 PE=1 SV=3
Collagen alpha−1(IV) chain OS=Mus musculus GN=Col4a1 PE=1 SV=1
Collagen alpha−2(IV) chain OS=Mus musculus GN=Col4a2 PE=1 SV=4
Heat shock cognate 71 kDa protein OS=Mus musculus GN=Hspa8 PE=1 SV=1
Fructose−bisphosphate aldolase A OS=Mus musculus GN=Aldoa PE=1 SV=2
SPARC OS=Mus musculus GN=Sparc PE=1 SV=1
Filamin, alpha OS=Mus musculus GN=Flna PE=1 SV=1
Insulin−like growth factor−binding protein 7 OS=Mus musculus GN=Igfbp7 PE=1 SV=1
Extracellular matrix protein 1 OS=Mus musculus GN=Ecm1 PE=1 SV=2
Thrombospondin 1 OS=Mus musculus GN=Thbs1 PE=1 SV=1
Prothymosin alpha OS=Mus musculus GN=Ptma PE=1 SV=2
Laminin subunit alpha−4 OS=Mus musculus GN=Lama4 PE=1 SV=2
Laminin subunit alpha−5 OS=Mus musculus GN=Lama5 PE=1 SV=4
Follistatin−related protein 1 OS=Mus musculus GN=Fstl1 PE=1 SV=2
Clusterin OS=Mus musculus GN=Clu PE=1 SV=1
Carboxypeptidase E OS=Mus musculus GN=Cpe PE=1 SV=2
Nidogen−1 OS=Mus musculus GN=Nid1 PE=1 SV=2
Cationic trypsin OS=Bos taurus PE=1 SV=3
Myosin−9 OS=Mus musculus GN=Myh9 PE=1 SV=4
Nucleobindin−1 OS=Mus musculus GN=Nucb1 PE=1 SV=2
Versican core protein OS=Mus musculus GN=Vcan PE=1 SV=1
Ceruloplasmin OS=Mus musculus GN=Cp PE=1 SV=1
Elongation factor 2 OS=Mus musculus GN=Eef2 PE=1 SV=2
Phosphoglycerate kinase 1 OS=Mus musculus GN=Pgk1 PE=1 SV=4
Syndecan−4 OS=Mus musculus GN=Sdc4 PE=1 SV=1
Pyruvate kinase PKM OS=Mus musculus GN=Pkm PE=1 SV=4
Thrombospondin−2 OS=Mus musculus GN=Thbs2 PE=1 SV=2
AHNAK nucleoprotein (desmoyokin) OS=Mus musculus GN=Ahnak PE=1 SV=1
Annexin A1 OS=Mus musculus GN=Anxa1 PE=1 SV=2
Moesin OS=Mus musculus GN=Msn PE=1 SV=3
Amyloid beta A4 protein OS=Mus musculus GN=App PE=1 SV=3
Prelamin−A/C OS=Mus musculus GN=Lmna PE=1 SV=2
Tropomyosin alpha−4 chain OS=Mus musculus GN=Tpm4 PE=1 SV=3
Protein disulfide−isomerase A3 OS=Mus musculus GN=Pdia3 PE=1 SV=2
Peptidyl−prolyl cis−trans isomerase B OS=Mus musculus GN=Ppib PE=1 SV=2
Myristoylated alanine−rich C−kinase substrate OS=Mus musculus GN=Marcks PE=1 SV=2
Latent−transforming growth factor beta−binding protein 1 OS=Mus musculus GN=Ltbp1 PE=1 SV=2
14−3−3 protein epsilon OS=Mus musculus GN=Ywhae PE=1 SV=1
Ubiquitin−40S ribosomal protein S27a OS=Homo sapiens GN=RPS27A PE=1 SV=2
Caldesmon 1 OS=Mus musculus GN=Cald1 PE=1 SV=1
14−3−3 protein zeta/delta OS=Mus musculus GN=Ywhaz PE=1 SV=1
Nucleolin OS=Mus musculus GN=Ncl PE=1 SV=2
Collagen alpha−1(XVIII) chain OS=Mus musculus GN=Col18a1 PE=1 SV=1
Tropomyosin alpha−1 chain OS=Mus musculus GN=Tpm1 PE=1 SV=1
Protein disulfide−isomerase OS=Mus musculus GN=P4hb PE=1 SV=2
Insulin−like growth factor−binding protein 6 OS=Mus musculus GN=Igfbp6 PE=2 SV=2
Alpha actinin 1a OS=Mus musculus GN=Actn1 PE=1 SV=1
Endoplasmin OS=Mus musculus GN=Hsp90b1 PE=1 SV=2
Fibrillin−1 OS=Mus musculus GN=Fbn1 PE=1 SV=2
Metalloproteinase inhibitor 2 OS=Mus musculus GN=Timp2 PE=1 SV=2
Keratin, type II cytoskeletal 1 OS=Homo sapiens GN=KRT1 PE=1 SV=6
Elongation factor 1−alpha 1 OS=Mus musculus GN=Eef1a1 PE=1 SV=3
Alpha−enolase OS=Mus musculus GN=Eno1 PE=1 SV=3
78 kDa glucose−regulated protein OS=Mus musculus GN=Hspa5 PE=1 SV=3
Neutral alpha−glucosidase AB OS=Mus musculus GN=Ganab PE=1 SV=1
Gremlin−1 OS=Mus musculus GN=Grem1 PE=2 SV=1
Plectin OS=Mus musculus GN=Plec PE=1 SV=3
Connective tissue growth factor OS=Mus musculus GN=Ctgf PE=2 SV=3
Complement C3 OS=Mus musculus GN=C3 PE=1 SV=3
Low−density lipoprotein receptor OS=Mus musculus GN=Ldlr PE=1 SV=1
Mesothelin OS=Mus musculus GN=Msln PE=1 SV=1
Protein SET (Fragment) OS=Mus musculus GN=Set PE=1 SV=1
Peptidyl−prolyl cis−trans isomerase C OS=Mus musculus GN=Ppic PE=1 SV=1
Peroxiredoxin−1 (Fragment) OS=Mus musculus GN=Prdx1 PE=1 SV=8
Tubulin alpha−1B chain OS=Mus musculus GN=Tuba1b PE=1 SV=2
Out at first protein homolog OS=Mus musculus GN=Oaf PE=2 SV=1
Cathepsin L1 OS=Mus musculus GN=Ctsl PE=1 SV=2
Transgelin−2 OS=Mus musculus GN=Tagln2 PE=1 SV=4
Vinculin OS=Mus musculus GN=Vcl PE=1 SV=4
Sulfhydryl oxidase 1 OS=Mus musculus GN=Qsox1 PE=1 SV=1
Decorin OS=Mus musculus GN=Dcn PE=1 SV=1
Annexin (Fragment) OS=Mus musculus GN=Anxa2 PE=1 SV=1
Keratin, type I cytoskeletal 10 OS=Homo sapiens GN=KRT10 PE=1 SV=6
Nucleoside diphosphate kinase OS=Mus musculus GN=Gm20390 PE=3 SV=1
Serpin H1 OS=Mus musculus GN=Serpinh1 PE=1 SV=3
Protein CYR61 OS=Mus musculus GN=Cyr61 PE=1 SV=1
L−lactate dehydrogenase OS=Mus musculus GN=Ldha PE=1 SV=1
Filamin−B OS=Mus musculus GN=Flnb PE=1 SV=3
Tropomyosin alpha−3 chain OS=Mus musculus GN=Tpm3 PE=1 SV=1
Cathepsin B OS=Mus musculus GN=Ctsb PE=1 SV=2
DNA segment, Chr 6, Wayne State University 176, expressed, isoform CRA_a OS=Mus musculus GN=Fam3c PE=1 SV=1
Matrilin−2 OS=Mus musculus GN=Matn2 PE=1 SV=1
Transgelin OS=Mus musculus GN=Tagln PE=1 SV=3
Biglycan OS=Mus musculus GN=Bgn PE=1 SV=1
Cadherin−2 OS=Mus musculus GN=Cdh2 PE=1 SV=2
Transitional endoplasmic reticulum ATPase OS=Mus musculus GN=Vcp PE=1 SV=4
T−complex protein 1 subunit theta OS=Mus musculus GN=Cct8 PE=1 SV=3
Collagen alpha−1(XI) chain OS=Mus musculus GN=Col11a1 PE=1 SV=2
Proliferation−associated protein 2G4 OS=Mus musculus GN=Pa2g4 PE=1 SV=3
Immunoglobulin superfamily containing leucine−rich repeat protein OS=Mus musculus GN=Islr PE=1 SV=1
60S ribosomal protein L7 (Fragment) OS=Mus musculus GN=Rpl7 PE=1 SV=1
Heat shock protein HSP 90−alpha OS=Mus musculus GN=Hsp90aa1 PE=1 SV=4
14−3−3 protein theta (Fragment) OS=Mus musculus GN=Ywhaq PE=1 SV=1
Semaphorin−3C OS=Mus musculus GN=Sema3c PE=1 SV=2
Alpha−actinin−4 OS=Mus musculus GN=Actn4 PE=1 SV=1
Collagen alpha−1(II) chain OS=Mus musculus GN=Col2a1 PE=1 SV=2
Uncharacterized protein OS=Mus musculus GN=Rpl7a−ps3 PE=4 SV=1
Insulin−like growth factor−binding protein 4 OS=Mus musculus GN=Igfbp4 PE=1 SV=2
Transketolase OS=Mus musculus GN=Tkt PE=1 SV=1
Tropomyosin beta chain OS=Mus musculus GN=Tpm2 PE=1 SV=1
Exostosin−2 OS=Mus musculus GN=Ext2 PE=1 SV=2
Calsyntenin−1 OS=Mus musculus GN=Clstn1 PE=1 SV=1
Protein−lysine 6−oxidase OS=Mus musculus GN=Lox PE=1 SV=1
60S ribosomal protein L12 OS=Mus musculus GN=Rpl12 PE=1 SV=2
Platelet−activating factor acetylhydrolase OS=Mus musculus GN=Pla2g7 PE=2 SV=2
Syndecan−1 OS=Mus musculus GN=Sdc1 PE=1 SV=1
Malate dehydrogenase, cytoplasmic OS=Mus musculus GN=Mdh1 PE=1 SV=3
Peroxidasin homolog OS=Mus musculus GN=Pxdn PE=1 SV=1
40S ribosomal protein S9 OS=Mus musculus GN=Rps9 PE=1 SV=3
Elongation factor 1−delta OS=Mus musculus GN=Eef1d PE=1 SV=1
Lipoprotein lipase OS=Mus musculus GN=Lpl PE=1 SV=3
Histone H1.4 OS=Mus musculus GN=Hist1h1e PE=1 SV=2
40S ribosomal protein SA OS=Mus musculus GN=Rpsa PE=1 SV=4
40S ribosomal protein S3 OS=Mus musculus GN=Rps3 PE=1 SV=1
Rho GDP−dissociation inhibitor 1 OS=Mus musculus GN=Arhgdia PE=1 SV=3
Amyloid−like protein 2 OS=Mus musculus GN=Aplp2 PE=1 SV=4
Procollagen−lysine,2−oxoglutarate 5−dioxygenase 1 OS=Mus musculus GN=Plod1 PE=1 SV=1
Protein NOV homolog OS=Mus musculus GN=Nov PE=1 SV=1
Heat shock 70 kDa protein 4 OS=Mus musculus GN=Hspa4 PE=1 SV=1
Metalloproteinase inhibitor 1 OS=Mus musculus GN=Timp1 PE=1 SV=2
Heterogeneous nuclear ribonucleoprotein A/B OS=Mus musculus GN=Hnrnpab PE=1 SV=1
Adenosylhomocysteinase OS=Mus musculus GN=Ahcy PE=1 SV=3
Hsc70−interacting protein OS=Mus musculus GN=St13 PE=1 SV=1
Triosephosphate isomerase OS=Mus musculus GN=Tpi1 PE=1 SV=4
T−complex protein 1 subunit gamma OS=Mus musculus GN=Cct3 PE=1 SV=1
Heterogeneous nuclear ribonucleoprotein K OS=Mus musculus GN=Hnrnpk PE=1 SV=1
Tubulointerstitial nephritis antigen−like OS=Mus musculus GN=Tinagl1 PE=1 SV=1
Inter−alpha trypsin inhibitor, heavy chain 2 OS=Mus musculus GN=Itih2 PE=1 SV=1
Tubulin beta−5 chain OS=Mus musculus GN=Tubb5 PE=1 SV=1
Fibulin−1 OS=Mus musculus GN=Fbln1 PE=1 SV=2
45 kDa calcium−binding protein OS=Mus musculus GN=Sdf4 PE=1 SV=1
Proprotein convertase subtilisin/kexin type 9 OS=Mus musculus GN=Pcsk9 PE=1 SV=2
60S ribosomal protein L13 OS=Mus musculus GN=Rpl13 PE=1 SV=3
Rab GDP dissociation inhibitor beta OS=Mus musculus GN=Gdi2 PE=1 SV=1
Valine−−tRNA ligase (Fragment) OS=Mus musculus GN=Vars PE=1 SV=1
60S acidic ribosomal protein P2 OS=Mus musculus GN=Rplp2 PE=1 SV=3
Eukaryotic translation initiation factor 4B OS=Mus musculus GN=Eif4b PE=1 SV=1
Angiopoietin−related protein 4 OS=Mus musculus GN=Angptl4 PE=2 SV=1
60S ribosomal protein L4 OS=Mus musculus GN=Rpl4 PE=1 SV=3
High mobility group protein B1 OS=Mus musculus GN=Hmgb1 PE=1 SV=1
Receptor−type tyrosine−protein phosphatase kappa OS=Mus musculus GN=Ptprk PE=1 SV=1
Angiotensin−converting enzyme (Fragment) OS=Mus musculus GN=Ace PE=1 SV=1
Fascin OS=Mus musculus GN=Fscn1 PE=1 SV=4
Heterogeneous nuclear ribonucleoproteins A2/B1 OS=Mus musculus GN=Hnrnpa2b1 PE=1 SV=2
Protein disulfide−isomerase A6 OS=Mus musculus GN=Pdia6 PE=1 SV=1
Myosin regulatory light chain 12B OS=Mus musculus GN=Myl12b PE=1 SV=2
Malate dehydrogenase, mitochondrial OS=Mus musculus GN=Mdh2 PE=1 SV=3
Golgi apparatus protein 1 (Fragment) OS=Mus musculus GN=Glg1 PE=1 SV=1
40S ribosomal protein S8 OS=Mus musculus GN=Rps8 PE=1 SV=2
Neural cell adhesion molecule 1 OS=Mus musculus GN=Ncam1 PE=1 SV=1
Proliferating cell nuclear antigen OS=Mus musculus GN=Pcna−ps2 PE=3 SV=1
Keratin, type II cytoskeletal 2 epidermal OS=Homo sapiens GN=KRT2 PE=1 SV=2
Glucosidase 2 subunit beta OS=Mus musculus GN=Prkcsh PE=1 SV=1
Latent−transforming growth factor beta−binding protein 3 OS=Mus musculus GN=Ltbp3 PE=1 SV=1
40S ribosomal protein S11 OS=Mus musculus GN=Rps11 PE=1 SV=1
Adenylosuccinate synthetase isozyme 2 OS=Mus musculus GN=Adss PE=1 SV=2
Importin−5 OS=Mus musculus GN=Ipo5 PE=1 SV=3
−2
−1
0
1
2

**Figure S7**

**A.**

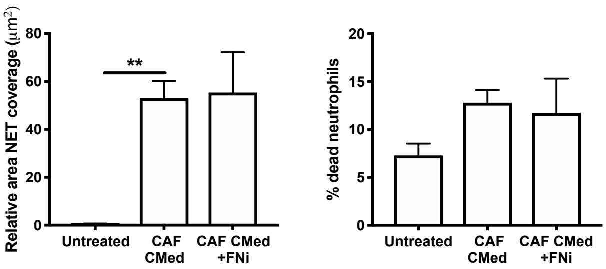


**B.**

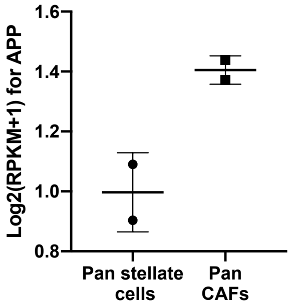


**C.**

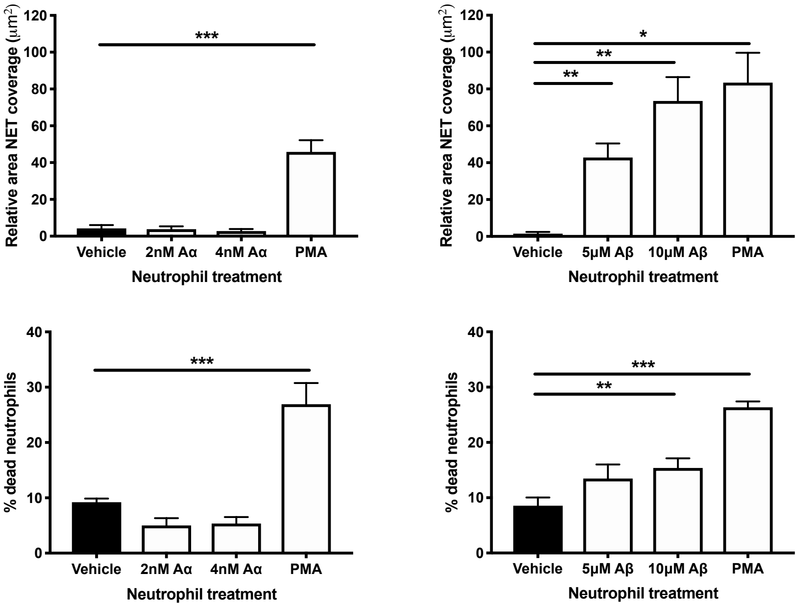

#### Figure S8

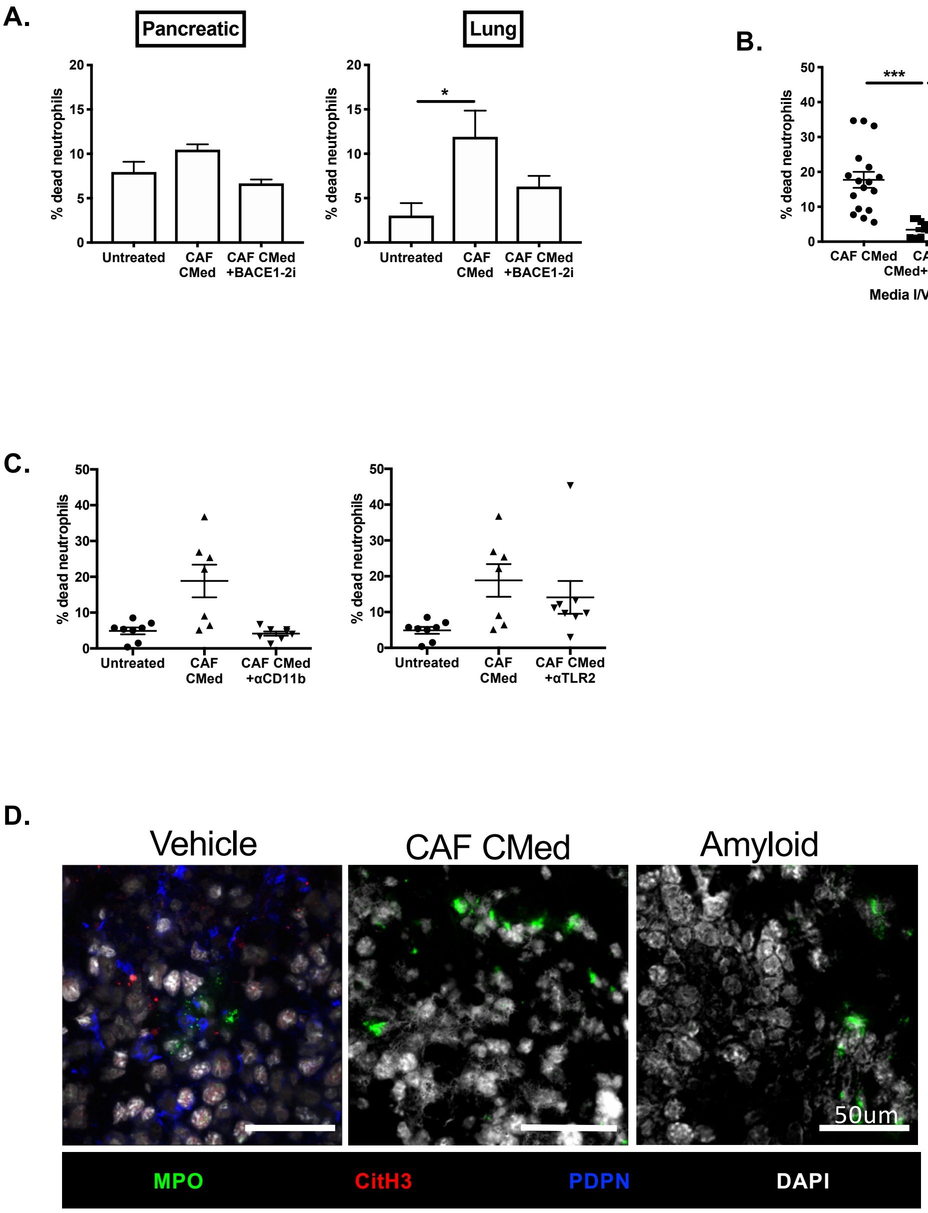
A.
Pancreatic
Lung
% dead neutrophils
Untreated
CAF CMed
CAF CMed +BACE1-2i
*
B.
% dead neutrophils
***
***
CAF CMed
CAF CMed+BACEi
rAmyloid β
Media I/V infused
C.
% dead neutrophils
Untreated
CAF CMed
CAF CMed +αCD11b
CAF CMed +αTLR2
D.
Vehicle
CAF CMed
Amyloid
50um
MPO
CitH3
PDPN
DAPI

#### Figure S9

A.

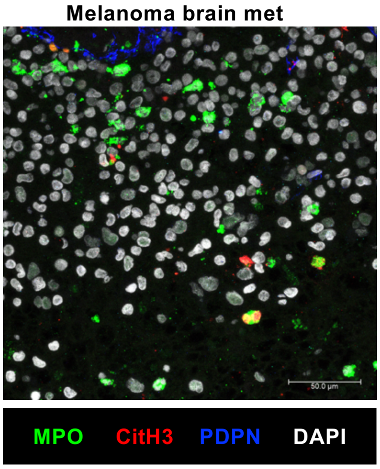
Melanoma brain met
50.0 µm
MPO
CitH3
PDPN
DAPI

B.

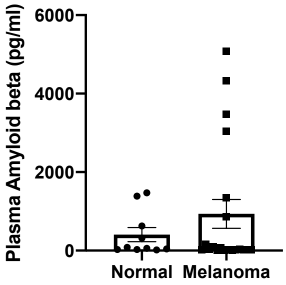
Plasma Amyloid beta (pg/ml)
6000
4000
2000
0
Normal
Melanoma
